# Dynamic nanoscale architecture of synaptic vesicle fusion in mouse hippocampal neurons

**DOI:** 10.1101/2025.02.11.635788

**Authors:** Jana Kroll, Uljana Kravčenko, Mohsen Sadeghi, Christoph A. Diebolder, Lia Ivanov, Małgorzata Lubas, Thiemo Sprink, Magdalena Schacherl, Mikhail Kudryashev, Christian Rosenmund

## Abstract

During neurotransmission, presynaptic action potentials trigger synaptic vesicle fusion with the plasma membrane within milliseconds. To visualize membrane dynamics before, during, and right after vesicle fusion at central synapses under near-native conditions, we developed an experimental strategy for time-resolved *in situ* cryo-electron tomography with millisecond temporal resolution. We coupled optogenetic stimulation with cryofixation and confirmed the stimulation-induced release of neurotransmitters via cryo-confocal microscopy of a fluorescent glutamate sensor. Our morphometric analysis of tomograms from stimulated and control synapses allowed us to characterize five states of vesicle fusion intermediates ranging from stalk formation to the formation, opening, and collapsing of a fusion pore. Based on these measurements, we generated a coarse-grained simulation of a synaptic vesicle approaching the active zone membrane. Both, our morphofunctional and computational analyses, support a model in which calcium-triggered fusion is initiated from synaptic vesicles in close proximity to the active zone membrane, whereby neither tight docking nor an induction of membrane curvature at the active zone are favorable. Numbers of filamentous tethers closely correlated to the distance between vesicle and membrane, but not to their respective fusion readiness, indicating that the formation of multiple tethers is required for synaptic vesicle recruitment preceding fusion.

## Main

Neuronal exocytosis is initiated by the recruitment of neurotransmitter-filled synaptic vesicles (SVs) to release sites at the active zone (AZ), where they are coupled to voltage-gated calcium channels and primed for fusion^1,2^. During SV fusion, the hydrophobic lipid bilayers of the SV and the AZ membrane are brought into close proximity and perturbed to enable the integration of the SV membrane into the cell membrane^3^. The synaptic fusion machinery, consisting of soluble N-ethylmaleimide-sensitive factor attachment protein receptors (SNAREs), regulatory proteins like Munc13 and Munc18, and the calcium sensor synaptotagmin-1, catalyzes the fusion reaction, helping to overcome energy barriers imposed by repulsive forces during SV-AZ membrane apposition and lipid reorganization^1,3,4^. Munc13 is not only required for bridging the cellular and vesicular membranes^5,6^ but also, together with Munc18, aids in the formation of SNARE complexes^7^. An SV has reached a primed state when trans-SNARE complexes have formed and closely interact with synaptotagmin-1 and complexin^8,9^. Calcium influx triggers membrane binding of synaptotagmin-1, accompanied by a rearrangement of the synaptotagmin-1-SNARE complex interface that in turn initiates fast neurotransmitter release by exerting work on the plasma membrane domains of SNARE proteins^10–12^.

Recent advances in electron microscopy (EM) fixation techniques have allowed for the visualization of SV fusion at small nerve terminals of central synapses. Using optogenetic or electrical stimulation and high-pressure freezing, followed by freeze substitution and EM (“flash-and-freeze” and “zap-and-freeze”, respectively), the activity-induced rearrangement of the coarse synaptic ultrastructure has been described^13–18^. Most fusion events, identified by small invaginations or pits at the AZ membrane, were observed 5-8 ms after action potential induction^13^, going along with a decreased number of morphologically docked SVs^13,14,16,17^. In addition to these and other studies using freeze-substituted and resin embedded EM samples, the synaptic architecture of isolated synaptosomes, cultured neurons, and even brain slices has recently been characterized using *in situ* cryo-electron tomography (cryo-ET), which allows for a much higher structurally interpretable resolution and near-native sample preservation^19–23^. These cryo-ET studies showed that most SVs are linked to each other via pleomorphic interconnectors and connected to the AZ membrane via filamentous tethers^19,24–27^, whereby the formation of multiple tethers was suggested to form a prerequisite for SV priming and fusion^25,26^. Not only Munc13^19,24,27^, but also SNARE proteins are suggested to be involved in the (dynamic) formation of these tethers^19,26–28^.

While *in situ* cryo-ET enables a molecular-level visualization of cellular landscapes, adding millisecond (ms) temporal resolution has so far been challenging^29^. Although a small number of SV fusion intermediates have been captured using a spraying technique for synaptic stimulation recently^26^, a systematic characterization of SV fusion *in situ* is still lacking. Our current understanding of membrane dynamics during SV fusion is therefore essentially built on *in vitro*^30–32^ and *in silico*^11,33–36^ studies of membrane fusion events. However, it is hard to predict how well the experimentally or computationally designed conditions of these studies reflect the actual cellular environment, including not only the interactome of proteins and lipids, but also biophysical properties like membrane tension, lipid rafts or phase separation.

In this study, we developed a time-resolved *in situ* cryo-ET workflow to enable a comprehensive reexamination of SV fusion under near-physiological conditions and within the native cellular environment. We coupled optogenetic stimulation with plunge freezing of cultured hippocampal neurons and confirmed successful stimulation using cryo-confocal microscopy of the glutamate sensor iGluSnFR3. We characterized the dynamic nanoscale architecture of SV fusion intermediates and used this structural information for a coarse-grained simulation of SV fusion initiation. We further morphometrically analyzed the distribution and tethering of membrane-near SVs, revealing a stimulation-induced reduction of SVs within a distance of 6 nm to the AZ membrane and a correlation between SV distances and tether numbers.

## Coupled optogenetic stimulation and cryofixation of mouse hippocampal neurons

To characterize the synaptic nanoscale architecture during and shortly after neurotransmitter release, we developed a workflow combining optogenetics and *in situ* cryo-ET (**Fig. 1a**). For optogenetic stimulation, we expressed the channelrhodopsin-2 (ChR2) variant ChR2(E123T/T159C) in murine hippocampal neurons, which was shown to induce action potentials in a particularly fast and robust manner^37,38^. In addition to the established ChR2(E123T/T159C) version, which includes YFP for cellular localization (**Fig. 1b**), we created additional constructs harboring a Cerulean or mScarlet to avoid potential spectral overlap with fluorescent sensors in subsequent experiments (**Suppl. Fig. 1a**). We performed electrophysiological recordings of neurons infected with ChR2(E123T/T159C)-YFP or one of the two new constructs ChR2(E123T/T159C)-Cerulean and ChR2(E123T/T159C)-mScarlet to assess their efficacy to induce action potentials. Onset of a light pulse resulted reliably in an action potential with an average delay of 4.6-4.8 ms, a similar value as described for ChR2(E123T/T159C)-YFP in a similar experimental setting^18^ (**Suppl. Fig. 1b**), and a robust response to light stimuli up to a frequency of 40 Hz for all three tested constructs (**Suppl. Fig. 1c**).

**Fig. 1:**
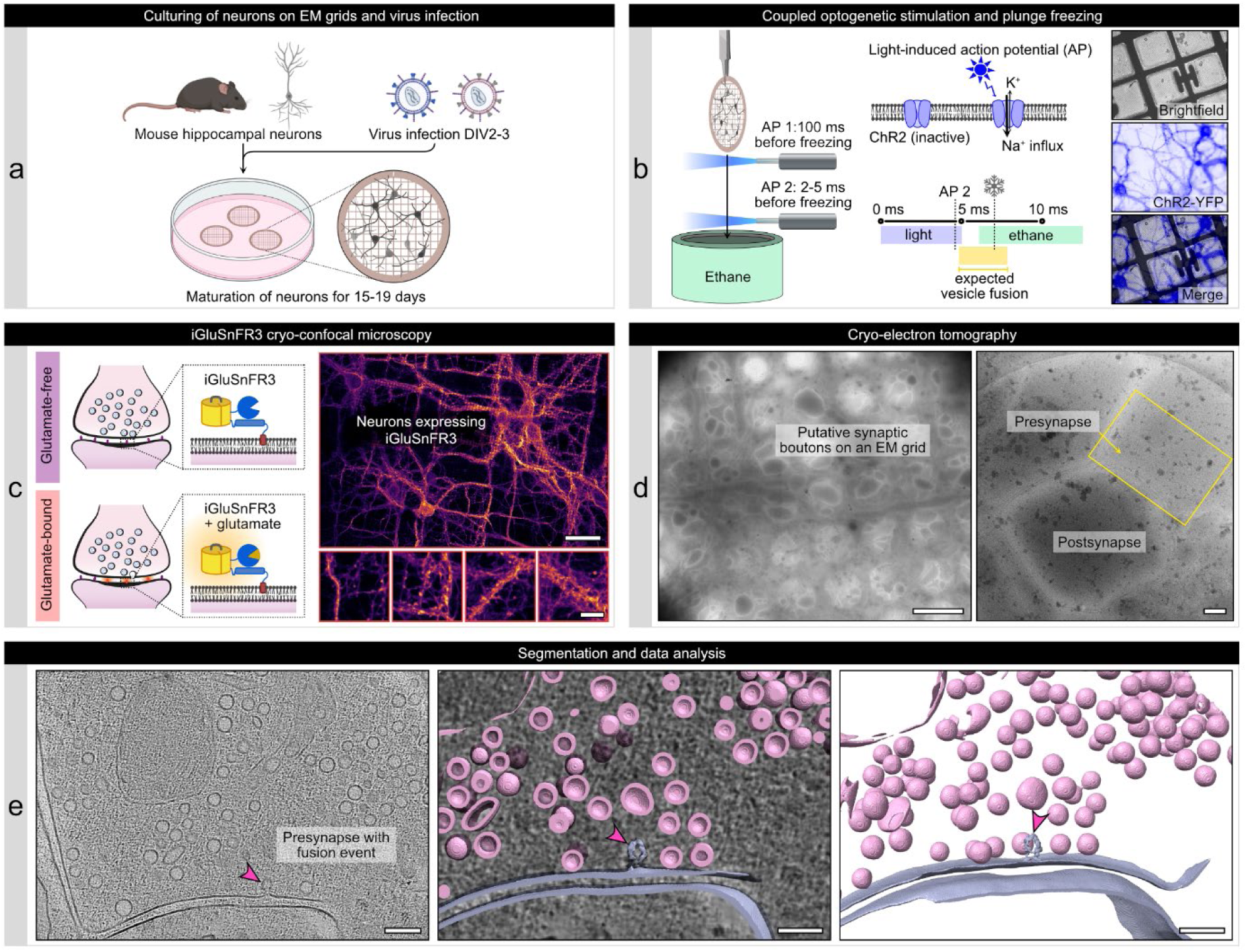
Workflow combining optogenetic stimulation of neurons, iGluSnFR cryo-confocal microscopy, and *in situ* cryo-ET. (**a**) Mouse hippocampal neurons are cultured on EM grids and infected with viruses for ChR2(ET/TC) and iGluSnFR3 expression. (**b**) Light pulses induce action potentials 100 ms (stim1) and 2-3 ms (stim2) before cryofixation of neurons via plunge freezing. Right panels: Cryo-fluorescence microscopy of ChR2(ET/TC)- YFP in plunge frozen neurons. (**c**). The upper right panel shows an overview of several neurons on an EM grid, scale bar 50 µm, the lower panels are zoom-ins to individual neurites containing synapses, scale bar 10 µm. (**d**) Cryo-ET tilt series were acquired from stimulated and control EM grids. Left: overview of a grid mesh with neurites and synaptic boutons, scale bar 5 µm. Right: Synapse within a hole of a holey carbon grid, scale bar 200 nm. (**e**) Tomograms are reconstructed from tilt series and used for segmentation and data analysis. The tomogram slice and segmentation show a stimulated synapse with ongoing SV fusion (pink arrowhead). Scale bars 200 nm. Schematic illustrations in panel a were generated with BioRender.

Neurons cultured on EM grids (for details of our cell culture setup, see **Suppl. Fig. 2**) were plunge frozen at DIV16-18 using a modified Vitrobot Mark IV (Thermo Fisher Scientific) plunge freezer equipped with an LED connected to optical fibers inside and below the chamber (**Fig. 1b** and **Suppl. Fig. 3a**). Optogenetic stimulation (2 pulses at 10 Hz) was performed at ∼37°C and an elevated extracellular calcium concentration of 4 mM to increase the vesicular release probability. The first stimulus was applied within the chamber at a maximum of 100 ms before vitrification and the second stimulus while the sample grid traveled towards the cooled ethane. The second stimulus started approximately 7 ms before vitrification, inducing an action potential 2-5 ms before the grid was dipped into cooled ethane (additional cooling time of the sample grid to 0°C < 1 ms^39^). The exact timing of the LED pulses and freezing were monitored using a high-speed camera (**Suppl. Fig. 3b**). Considering that most action potentials were induced 3-6 ms after light onset in our electrophysiological experiments and that the delay between action potential generation at the presynapse and synchronous neurotransmitter release is typically 1 ms or shorter^40^, our setup was well suited for cryofixing neurons shortly before, during, and directly after neurotransmitter release.

## Confirmation of neurotransmitter release using the glutamate sensor iGluSnFR3 in cryofixed neurons

After plunge freezing, we aimed to validate successful stimulation using a fluorescent biosensor for synaptic activity. For this purpose, we first characterized the kinetics and fluorescence intensity changes of the calcium sensor, SynGCaMP6f^41^, and different variants of the glutamate sensor, iGluSnFR, via live imaging of neurons cultured on coverslips (**Fig. 2a** and **Suppl. Fig. 4a**). Of all tested constructs, the fluorescent glutamate sensor iGluSnFR3.v857.GPI containing a GPI anchor for postsynaptic enrichment (^42^, from now iGluSnFR3) yielded the best-fitting properties with a maximum fluorescence intensity of 0.4 ± 0.03 ΔF/F_0_, (**Suppl. Fig. 4b**), an increase to half-maximum of τ_50%_ = 22.6 ± 3 ms (**Suppl. Fig. 4d**), and an increase to maximal intensity of τ_max_ = 64.7 ± 6.2 ms (**Suppl. Fig. 4c**). To examine the cellular localization of iGluSnFR3 signals, we performed a *post hoc* immunofluorescence staining of synaptic proteins on samples used for live imaging (**Suppl. Fig. 4f**). With this correlation, we could verify that action potential-induced iGluSnFR3 signals overlap primarily with the postsynaptic marker Homer1.

**Fig. 2:**
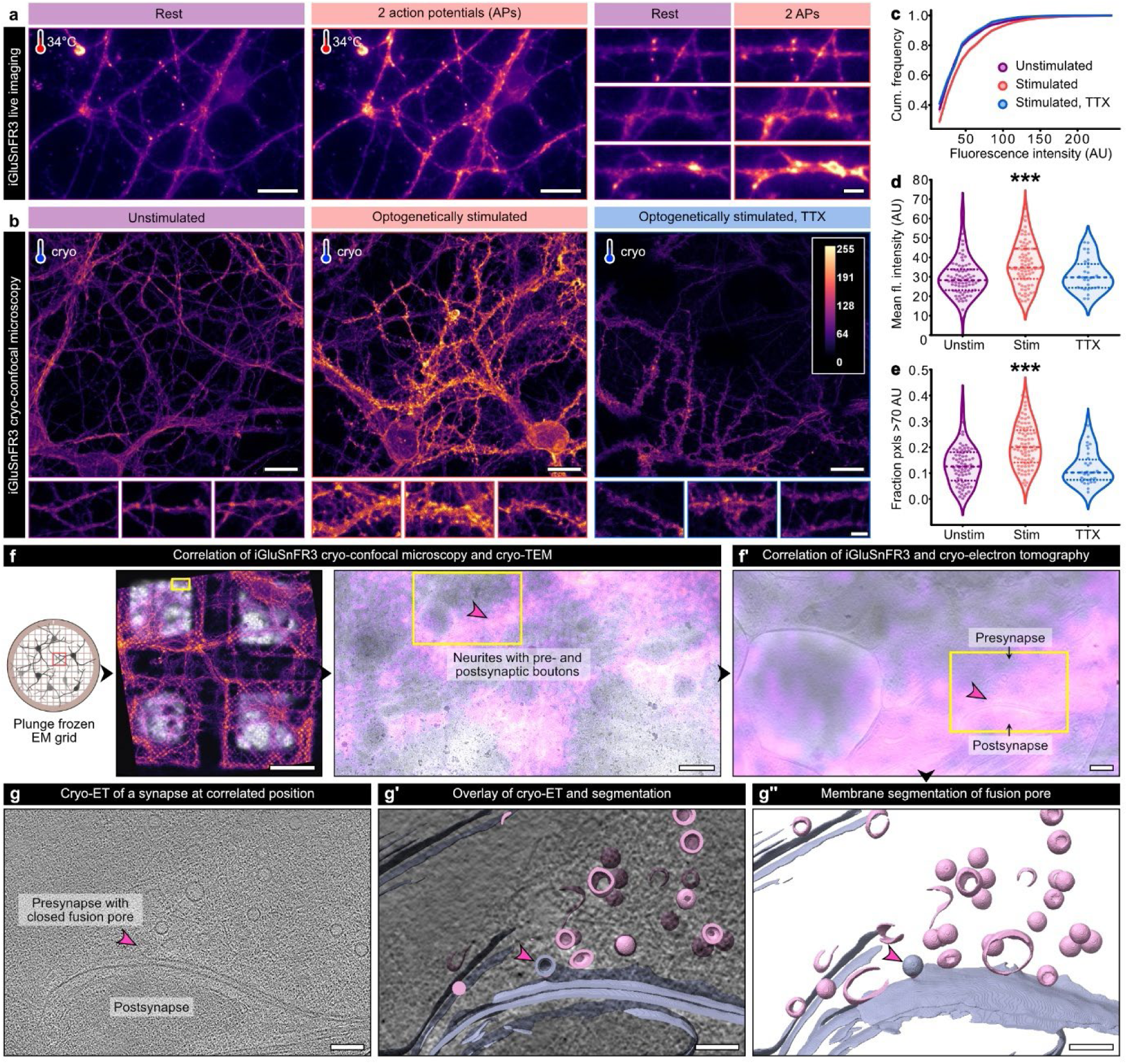
Confirmation of synaptic glutamate release in stimulated, cryofixed neurons. (**a**) Electrical field stimulation and live imaging of iGluSnFR3 at near-physiological temperature. Left panels: before stimulation, right panels: after stimulation. Scale bars: overviews 20 µm, zoom-ins 10 µm. (**b**) Maximum intensity projections of cryo-confocal stacks from unstimulated and optogenetically stimulated hippocampal neurons without and with TTX treatment. Scale bars upper panels: 20 µm, lower panels: 5 µm. (**c-e**) Cumulated fluorescence intensity histograms (c), mean fluorescence intensity (d) and fractions of pixels with a high fluorescence intensity (>70 AU) measured in areas containing individual neurites. unstimulated: N = 77 confocal stacks from 9 grids and 4 independent cultures; stimulated: N = 80 stacks from 13 grids and 4 cultures; stimulated TTX: N = 30 stacks from 3 grids and 2 cultures. Dashed lines in the violin plots indicate the median, dotted lines the 25% and 75% percentile. *** p < 0.001. (**f**) Correlative iGluSnFR3 cryo-confocal microscopy and cryo-TEM of a stimulated grid. Left panel: overview of four grid meshes, scale bar 50 µm. Right panel: zoom into a region containing synaptic boutons, scale bar 500 nm. (**f’**) Correlation of iGluSnFR3 fluorescence and a reconstructed tomogram slice of a synapse, scale bar 200 nm. (**g-g’**’) Tomogram slice (g), overlay (g’), and segmentation (g’’) of a putative closed fusion pore (pink arrowhead) from the correlated synapse in (f). Scale bars 100 nm. The schematic illustration in f was generated with BioRender.

Assuming that the glutamate-bound, highly-fluorescent conformation of iGluSnFR3 can be preserved under cryogenic conditions, we acquired cryo-confocal stacks of optogenetically stimulated and plunge frozen neurons expressing iGluSnFR3. We compared fluorescence intensities in neurites of unstimulated neurons, optogenetically stimulated neurons, and optogenetically stimulated and tetrodotoxin-treated (TTX, pharmacologically blocks sodium channels required for action potential induction) neurons of four independent cultures (two for TTX, **Fig. 2b**). In all four cultures, of which two were infected with ChR2(E123T/T159C)-YFP, one with ChR2(E123T/T159C)-mScarlet, and one culture with both, the mean fluorescence intensity of the stimulated samples was consistently higher than of the unstimulated samples, indicating that our setup combining optogenetic stimulation and plunge freezing works reliably.

We therefore pooled the measurements of the individual cultures and calculated fluorescence intensity histograms for all three conditions (**Fig. 2c**). In the stimulated samples, the mean fluorescence intensity of 36.3 ± 1.3 AU was significantly higher than in the two control conditions (unstimulated: 30.0 ± 1.1 AU, TTX-treated: 31.0 ± 1.5 AU, Kruskal-Wallis- test p < 0.001, **Fig. 2d**). Likewise, the fraction of pixels with a high fluorescence intensity (> 70 AU), likely reflecting glutamate-bound iGluSnFR3, was significantly increased after stimulation (20.6 ± 0.9% vs. 13.0 ± 0.9% and 11.9 ± 1.2% without stimulation and after TTX treatment, respectively, Kruskal-Wallis- test p < 0.001, **Fig. 2e**). Of note, the difference in mean fluorescence intensities of optogenetically stimulated and control conditions was less pronounced under cryogenic conditions than during live imaging, likely because low-expressing and non-responding neurons were excluded during live imaging, whereas cryo-confocal stacks were acquired from regions selected blindly.

To test if high iGluSnFR3 fluorescence intensity can be used as a marker for SV fusion events, we performed correlative cryo-confocal microscopy and cryo-EM (cryo-CLEM) on a stimulated EM grid (**Fig. 2f** and **Suppl. Fig. 5**). As visible in the overview of four grid meshes, the overall morphology of neurons was preserved in our on-grid cell culture system. For tilt series acquisition, we selected regions containing a high density of neurites but no cell somata (yellow box in the right panel of **Fig. 2f**). Correlating fluorescence and transmission electron microscopy (TEM), we observed the highest fluorescence intensity around large boutons likely resembling synapses (yellow arrowheads and box in **Suppl. Fig. 5**). The correlation of iGluSnFR3 fluorescence and tomogram slice (**Fig. 2f’**) revealed that the highest fluorescence intensity was visible at the synaptic cleft between a presynapse and a postsynaptic bouton. At the presynaptic AZ membrane, a putative forming fusion pore was observed (**Fig. 2g**).

## *In situ* cryo-ET of SV fusion intermediates

Having confirmed that neurons cultured on EM grids were optogenetically stimulated before being subsequently plunge frozen, we acquired cryo-ET data from those grids to morphometrically and biophysically characterize SV fusion. Although we could show that the iGluSnFR3 fluorescence signal is *per se* suited to select regions of interest for cryo-ET, we acquired tilt series without correlation of each position to avoid ice contaminations, which form during prolonged cryo-confocal microscopy, and to avoid the thinning of our samples via focused ion beam (FIB)-milling, going along with a lower sample throughput. We therefore acquired tilt series from regions with good cell and ice quality after confirming successful stimulation via cryo-fluorescence microscopy at only a few positions of the grids. From the acquired tilt series, we reconstructed 312 tomograms from stimulated and 95 tomograms from control (TTX-treated) samples. We screened each tomogram manually for synapses with a visible, cross-sectioned AZ. This resulted in 75 synapses in the stimulated and 28 synapses in the TTX-treated samples that were used for further analysis. Each AZ was then examined in more detail for membrane rearrangements that may be attributed to SV fusion (see **Suppl. Fig. 6** for examples of excluded structures). Based on all observed events, and in accordance with previously described SV fusion intermediates^3,25,26^, we defined seven categories: invagination of the AZ membrane (1), stalk formation (2), closed fusion pore (3), open fusion pore (4), dilating fusion pore (5), collapsing fusion pore (6), and small bumps (7) (**Fig. 3a, b, Suppl. Fig. 7,** see **Suppl. Table 1** for morphological criteria of each category). In addition to exemplary tomogram slices (**Fig. 3b**), we visualized 3D volumes of each category using UCSF ChimeraX^43^ (**Fig. 3c**). We further performed subtomogram averaging (StA) of selected fusion events from most categories using Dynamo^44^ (**Suppl. Fig. 7a**) and applied C61 symmetry (**Suppl. Fig. 7b**) to visualize the general membrane shape and bending. We were not able to generate an StA of open fusion pores because this category was particularly heterogeneous with open pore widths ranging from 2 to 18 nm.

**Fig. 3:**
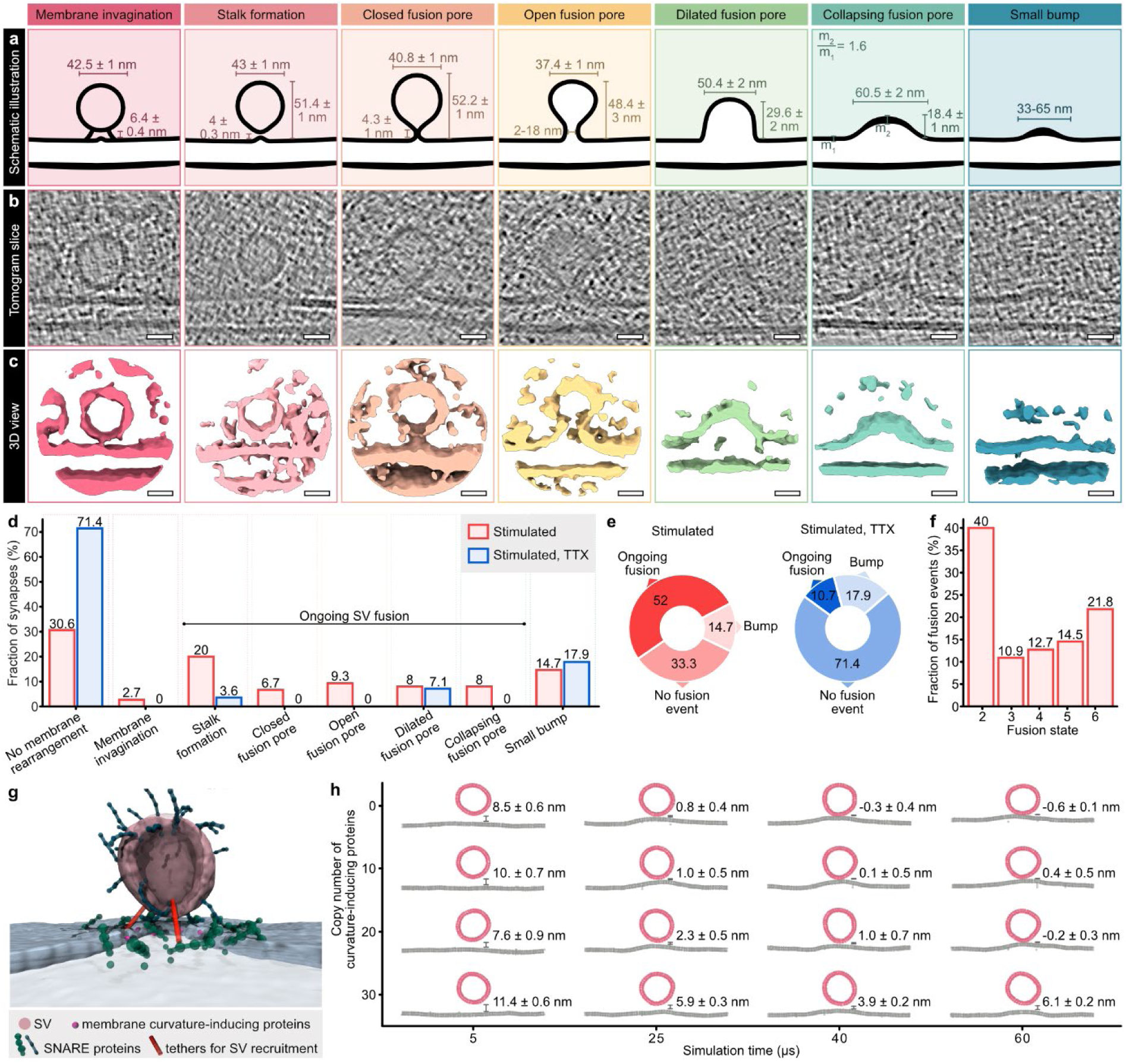
Cryo-ET of synaptic vesicle fusion states. (**a-c**) Schematic illustrations with details about size measurements (a), exemplary cryo-ET slices (b) and isosurfaces (c) of 7 categories of membrane rearrangements observed at synaptic active zones of stimulated neurons. Scare bars: 20 nm. (**d**) Fractions of synapses with or without membrane rearrangements in stimulated and stimulated, TTX-treated samples. (**e**) Fractions of synapses without membrane rearrangements, ongoing fusion, or bumps in stimulated and stimulated, TTX-treated samples. Synapses were counted as “ongoing fusion” if at least one stalk formation, closed, open, dilated, or collapsing fusion pore was observed. (**f**) Fractions of individual fusion states in stimulated synapses. Stimulated sample: N = 75 synapses from 3 grids and 2 independent freezings, TTX sample: N = 28 synapses from 1 grid. (**g**) Snapshot of a coarse-grained simulation of an SV approaching the active zone membrane, with particle-based representation of recruiting tethers, SNARE proteins and varying copy numbers of proteins inducing membrane curvature at the active zone below the SV. (**h**) Time evolution of the vertical distance between SV and active zone membrane, depending on the copy number of membrane curvature-inducing proteins.

SVs of category 1 (n = 10 SVs in stimulated synapses) were spherical and in a distance of 6.4 ± 0.4 nm to the AZ membrane (**Fig. 3a**), the AZ membrane below the center of the SV was slightly invaginated (**Suppl. Fig. 7c**). In category 2 (stalk formation, n = 22), SVs were droplet-shaped with an evagination at the SV bottom, while the AZ membrane was almost flat or slightly invaginated. At closed fusion pores belonging to category 3 (n = 6), the SV and AZ membrane had already fused, resulting in a continuous membrane. In comparison to category 2, closed fusion pores were slightly taller, the invagination of the AZ membrane was more pronounced. Open fusion pores (category 4, n = 7) were smaller than closed fusion pores, the width of the open pores was between 2.3 and 18.3 nm. Based on our definition, open pores contained outward (positive) and inward (negative) membrane curvature at the pore neck, whereas dilating pores (category 5, n = 8) only showed inward curvature at the junction of SV and cell membrane. The side walls of dilating pores were vertical or angular. Compared to dilating pores, collapsing fusion pores (category 6, n = 12) were lower and wider. Interestingly, the top membrane of collapsing fusion pores appeared thickened in relation to the surrounding AZ membrane by a factor of 1.6 (**Fig. 3a** and **Suppl. Fig. 7c**). Small bumps (category 7, n = 21) were again lower than collapsing fusion pores and varied in size and shape.

Compared to TTX-treated samples, we observed higher fractions of events in categories 1-6 in stimulated neurons (**Fig. 3d**), whereas the fraction of small bumps (7) was alike under both conditions. Closed and open fusion pores were only present in the stimulated sample without TTX treatment. Therefore, we defined membrane rearrangements of categories 2-6 as ongoing SV fusion and cell membrane invaginations below tethered, round SVs (category 1) as events likely preceding SV fusion. Based on this definition, we observed ongoing SV fusion in 52% (39/75 synapses), bumps in 14.7% (11/75), and no membrane rearrangements in 33.3% (25/75) of all stimulated synapses. In the TTX-treated group, we found fusion events in 10.7% (3/28 synapses), bumps in 17.9% (5/28), and no membrane rearrangements in 71.4% (20/28) of all synapses (**Fig. 3e**).

Since some synapses contained more than one fusion event, we further counted each fusion event individually (**Fig. 3f**). Of all fusion events observed in the stimulated samples (N = 55), the majority was stalk formation (40%), followed by collapsing fusion pores (21.8%). Closed (10.9%), open (12.7%), and dilating (14.5%) fusion pores were less prevalent. Presuming that more transient and volatile conditions are stochastically less likely to be captured during plunge freezing, our observed numbers of events per fusion state may serve as a morphological readout for their speed. Based on this assumption, fusing SVs may remain in state 2 (stalk formation) for comparatively longer, likely because energy barriers need to be overcome when the membranes of SV and AZ are approached and perturbed^3^. Alternatively or in addition, some of the formed stalks may not lead to SV fusion but instead get stuck or disassemble again^45,46^.

## Initiation of SV fusion via stalk formation

In previous studies, not only stalk formation but alternatively also (tight) docking has been suggested as prefusion state^3,31^. During tight docking, the SV approaches the AZ until the membranes are in direct and broad contact; the lipids of SV and AZ membrane are supposed to intermix until they reach a hemifusion (diaphragm) state. In our examples of stalk formation, the SVs were droplet-shaped and the average distance between SV and membrane was 4 ± 0.3 nm (smallest measured distance: 2.3 nm). This space between SV and AZ membrane appeared partially blurry in some of our examples (**Suppl. Fig. 7c**), which has previously been attributed to starting lipid intermixing^26^. In contrast, we observed one morphologically tightly docked SV at an AZ (**Suppl. Fig. 7d**), as well as one tightly docked SV (**Suppl. Fig. 7e**) and a putative hemifusion diaphragm (**Suppl. Fig. 7f**), both not in an AZ. Together, our observations indicate that a transition from tethering to stalk and fusion pore formation without (tight) docking is likely the predominant fusion mechanism.

Furthermore, a slight invagination of the AZ membrane (state 1) was present below 6.5% of all tethered SVs within a distance of 4-8 nm from the AZ membrane (10/155 SVs). We observed these invaginations at stimulated synapses with and without additional fusion events. To test whether an invagination of the AZ membrane is beneficial for SV fusion initiation and may thus precede stalk formation, we generated a coarse-grained simulation of an SV approaching the AZ membrane (**Fig. 3g**). In this model, proteins (e.g. resembling synaptotagmin-1) actively induce membrane curvature as soon as the SV has reached a distance of 6 nm to the AZ and also interact with SNAREs. We thereby incorporated size and distance measurements of our morphometric analyses (**Fig. 3a**, **Fig. 4c-g**, **Suppl. Fig. 9**). With this model, we tested the effects of different concentrations of membrane curvature-inducing proteins on SV approximation and SNARE complex formation. Interestingly, higher copy numbers of these proteins did not facilitate but rather impeded the recruitment of the SV, likely because SNARE complexes could not be formed efficiently anymore (**Fig. 3h**, **Suppl. Fig. 8**). Instead, 0 or 10 copies resulted in a fast SV approximation (in our model, SV fusion was not enabled). This means that although a slight invagination of the AZ membrane, as observed by us and in a previous study^26^, may precede SV fusion, it is unlikely induced by proteins like synaptotagmin- 1 but rather a consequence of SNARE zippering^10,12^.

**Fig. 4.**
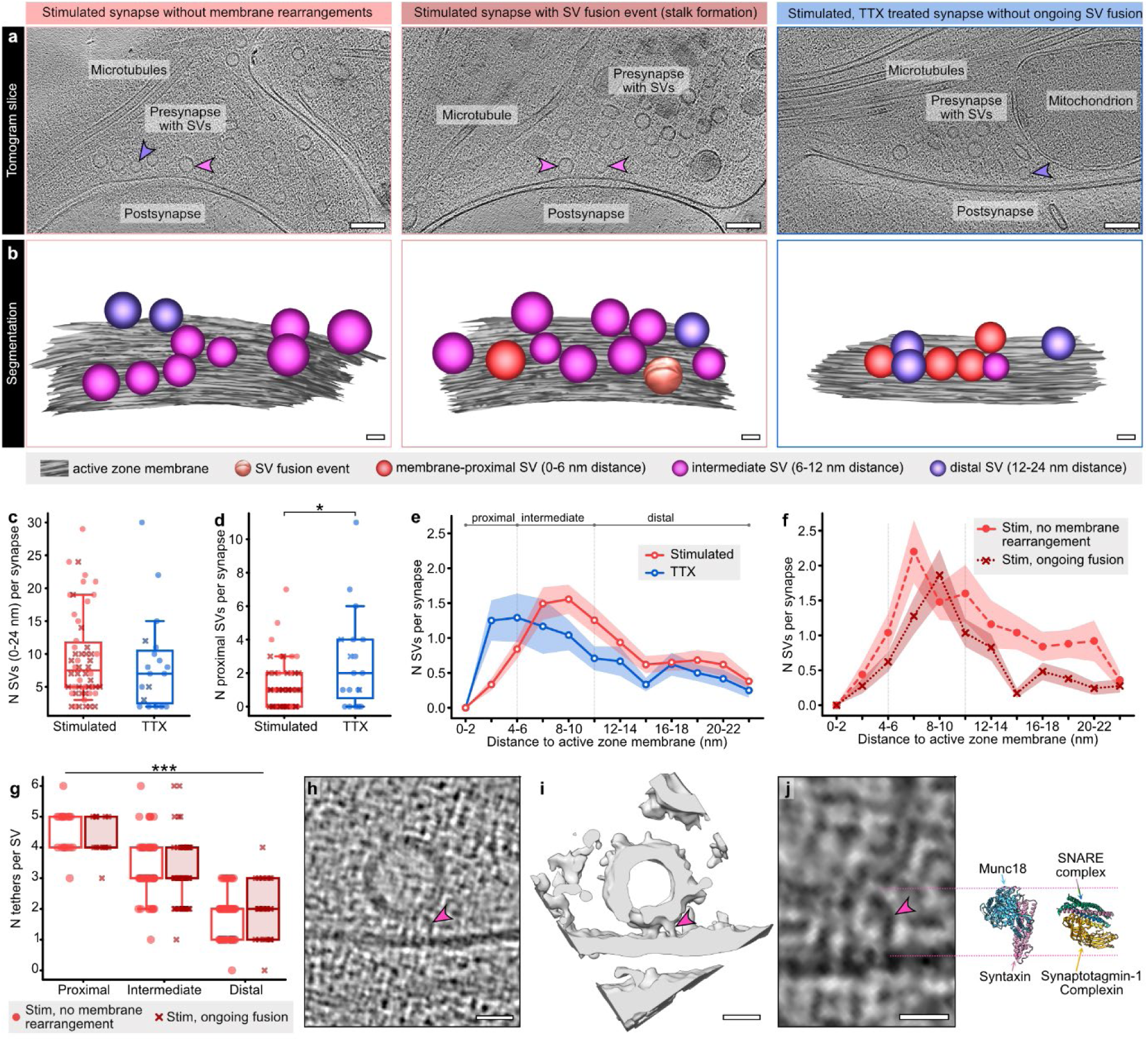
Stimulation-induced changes in the distribution and tethering of membrane-near SVs. (**a**) Exemplary cryo-ET slices of synapses without and with ongoing fusion in stimulated and stimulated, TTX-treated neurons. Scale bars 100 nm. (**b**) Manual segmentations of active zones and membrane-near SVs. Scale bars 20 nm. (**c, d**) Numbers of SVs per synapse with a max. distance of 24 nm (c) or 6 nm (proximal SVs, d). stim: N = 54 synapses, TTX: N = 19 synapses, * p < 0.05. (**e**) Distribution of membrane-near SVs in stimulated and stimulated, TTX-treated neurons. (**f**) Distribution of membrane-near SVs in synapses of stimulated neurons with ongoing fusion and without membrane rearrangements. No membrane rearrangement: N = 25 synapses, ongoing fusion: N = 29 synapses. (**g**) Numbers of tethers for membrane-proximal, intermediate (6-12 nm) and distal (12-24 nm) SVs of stimulated neurons. No membrane rearrangement: N = 145 SVs from 15 synapses, ongoing fusion: N = 114 SVs from 18 synapses, *** p < 0.001. (**h**) Exemplary tomogram slice of a multi-tethered SV. The pink arrowhead indicates one of the tethers. Scale bar 20 nm. (**i**) Isosurface of the same tethered SV. Scale bar 20 nm. (**j**) Atomic models of syntaxin/Munc18 (left, PDB: 4JEU) and an assembled SNARE complex with synaptotagmin-1 and complexin (right, PDB: 5W5D) in comparison to the tether indicated in (h) for size estimation, scale bar 10 nm. Light blue: Munc18, pink: syntaxin-1, green: SNAP-25, dark blue: synaptobrevin-2, yellow: complexin, orange: synaptotagmin-1.

## Depletion of membrane-proximal SVs in stimulated synapses

In addition to ongoing SV fusion events, we analyzed the distribution of tethered SVs within a distance of 24 nm from the AZ membrane (**Fig. 4a, 4b**). In previous EM studies, changes in the abundance of membrane-near SVs have been used as confirmation for successful action potential induction and subsequent release^13–18^. To test whether we could reproduce these findings with our workflow, we first compared all stimulated synapses (n = 515 SVs from 54 synapses) to TTX-treated synapses (n = 162 SVs, 19 synapses), whereby synapses with only bumps (state 7) were not included. While the total number of SVs within a distance of 24 nm was not significantly different between the two groups (Mann-Whitney test p = 0.477, **Fig. 4c**), we counted on average 1.4 ± 0.6 fewer SVs per AZ within a maximum distance of 6 nm (Mann-Whitney test p = 0.035, proximal SVs, **Fig. 4d, 4e**) in stimulated synapses. Between 6 and 12 nm (intermediate SVs), we observed slightly more SVs in stimulated synapses (**Suppl. Fig. 9c**). A comparable redistribution of SVs was reported using “zap-and-freeze” and freeze substitution, however, the effects were more drastic^13^.

In addition, we analyzed SV distributions of stimulated synapses with ongoing SV fusion (n = 29) and without membrane rearrangements (n = 25) individually. In the membrane-proximal SV pool (0-6 nm distance), we observed 0.6 ± 0.4 fewer SVs per AZ in stimulated synapses containing at least one fusion event (**Fig. 4f**). Consequently, also the subgroup of stimulated synapses without observed ongoing SV fusion had on average less membrane-proximal SVs than TTX-treated synapses (counted difference 1.1 ± 0.7 SVs, **Suppl. Fig. 9b, 9e**). It is likely that this subgroup of stimulated synapses without membrane rearrangements consists of synapses postfusion (neurotransmitter release has already taken place) and synapses without neurotransmitter release (non-releasing synapses). Assuming that the distribution of membrane-proximal SVs in postfusion synapses is comparable to synapses with ongoing fusion, whereas non-releasing synapses assumingly resemble TTX-treated synapses, we calculated the theoretical fraction of non-releasing synapses (see **Supplementary Methods**): Within the group of synapses without membrane rearrangements, the fraction of non-releasing synapses would be 17%. Of all stimulated synapses, the fraction of non-releasing synapses would be 8%, resulting in a theoretical synaptic release probability of 92% with our workflow. The release probability of excitatory hippocampal synapses at a calcium concentration of 4 mM and near-physiological temperature was reported to be ∼85%^47^. Beyond that, it is conceivable that SV replenishment starts already less than 11 ms after fusion^13^, potentially resulting in slightly more membrane-proximal SVs in the postfusion state than during fusion. In other words, we observed on average fewer membrane-proximal SVs in the subgroup of stimulated synapses without ongoing fusion than expected. Kusick and colleagues^13^ came to the same conclusion and attributed the low SV number to transient SV undocking during or shortly after fusion.

Previous work has shown that not only the distribution of SVs, but also the number of tethers dynamically changes during synaptic activity, whereby the formation of three or more tethers connecting SV and AZ membrane was suggested to be a morphological correlate of priming and a prerequisite for SV fusion^25,26^. Although our tomograms of synapses were comparatively thick, going along with a potentially worse signal-to-noise ratio than FIB-milled samples or purified synaptosomes, we were able to quantify tethers in our stimulated samples. Overall, we manually quantified tethers at 114 SVs from 18 synapses with ongoing fusion and 145 SVs from 15 synapses without membrane rearrangements. In both groups, we observed a linear correlation of tether number and distance between SV and AZ membrane (**Suppl. Fig. 9h**), whereby membrane-proximal SVs had the highest average number of tethers (4.6 ± 0.1 and 4.4 ± 0.2 for synapses without and with fusion event, respectively) and distal SVs the lowest (**Fig. 4g**). The number of tethers below SVs with slight membrane invagination was 4.2 ± 0.3 and did thus not differ from the other SVs within the same distance to the cell membrane. Of note, we observed short, vertical tethers predominantly below SVs (**Fig. 4h, Suppl. Fig. 10**) and longer, angular and curved tethers predominantly at the sides of membrane-proximal SVs (**Suppl. Fig. 10h**). For size estimation, we positioned atomic models of different exocytic proteins/complexes next to a tethered SV (**Fig. 4j**). For comparison with recent cryo-ET studies of FIB-milled synapses or synaptosomes^19,27^, we fitted these atomic models and corresponding density maps into 3D volumes of tethers connecting an SV to the AZ membrane (**Suppl. Fig. 10c-f**). Based on these fits, SNARE proteins may be involved in the formation of the short tethers observed here, as suggested previously^19,26,27^. However, not only assembled SNARE complexes together with synaptotagmin-1 and complexin (PDB: 5W5D^9^), which would indicate a primed state of the SV, are likely candidates. Syntaxin-1 and Munc18 (PDB: 4JEU^48^) or syntaxin-1, synaptobrevin-2 and Munc18 (PDB: 7UDB^49^), both representing states preceding SV priming, would fit equally well. Size-wise, Munc13 (PDB: 7T7V or 7T7X^50^) could be involved in the formation of longer angled tethers (**Suppl. Fig. 10h**), however, we cannot rule out that Munc13 is also part of short vertical tethers (**Suppl. Fig. 10f**). Of note, we also observed filaments connected to SV fusion intermediates, e.g. around the space between SV and AZ membrane during stalk formation (**Suppl. Fig. 7c**). Whether these filaments resemble parts of the SV fusion machinery, e.g. assembled SNARE complexes, needs to be investigated further.

## Multivesicular release and release site refilling

In 23% of all stimulated synapses with ongoing SV fusion (9/39 synapses), we observed multiple fusion events (**Fig. 5a, 5c**), whereby most of these synapses contained two (**Fig. 5d**). Likewise, we observed multiple small bumps per synapse in 55% of synapses containing bumps (6/11 synapses, **Fig. 5b, 5e**). At the synapses with MVR, the individual fusion events did not preferentially fall into the same category.

**Fig. 5:**
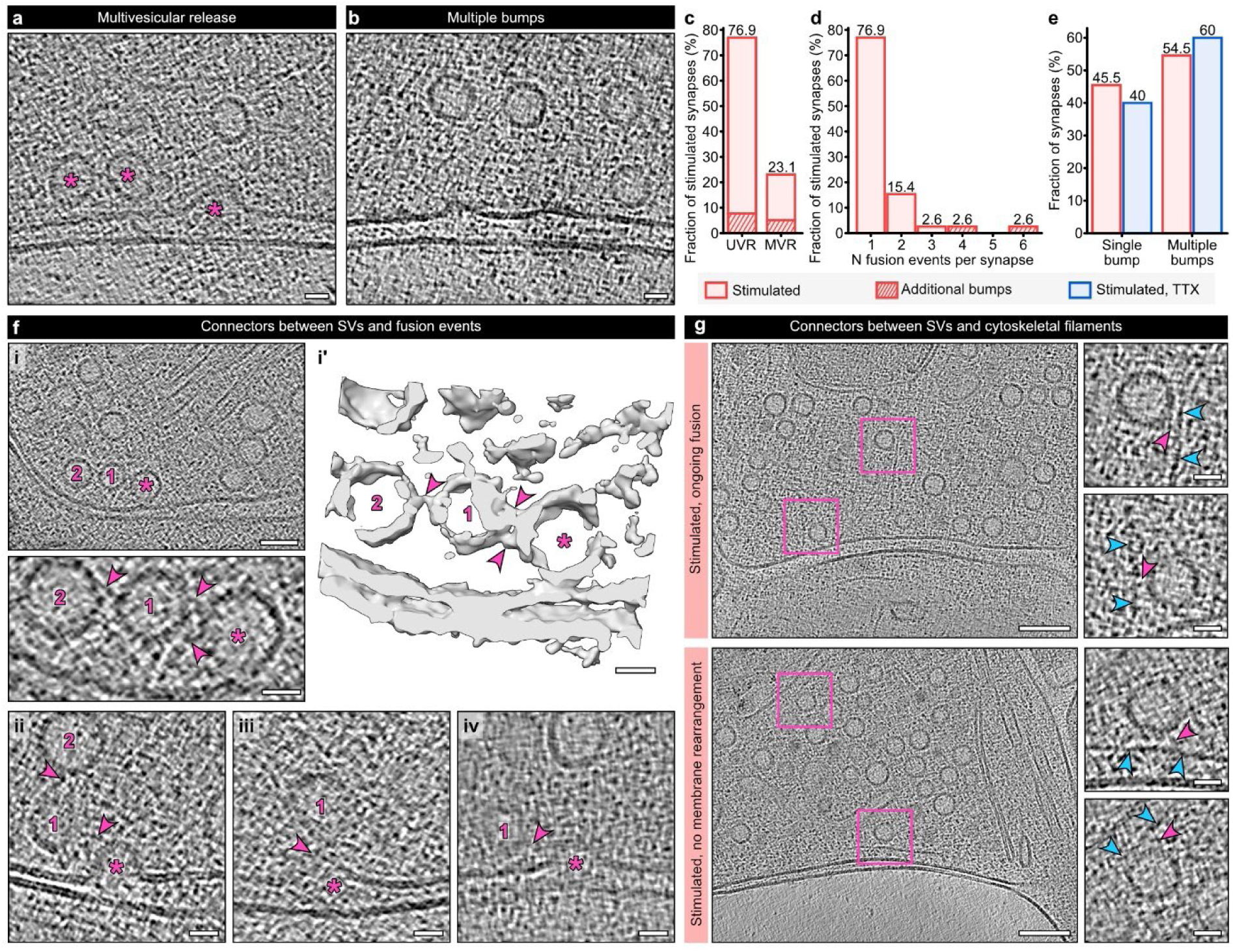
Multivesicular release and filamentous structures mediating SV resupply. (**a**) Exemplary tomogram slice of multivesicular release (MVR). The asterisks indicate two stalks and a dilated fusion pore at one active zone. Scale bar 20 nm. (**b**) Exemplary tomogram slice of multiple bumps. Scale bar 20 nm. (**c**) Fractions of synapses with univesicular release (UVR) and MVR in stimulated synapses. Synapses additionally containing bumps are indicated as shaded areas. (**d**) Numbers of fusion events per synapse in the stimulated sample. (**e**) Fractions of single and multiple bumps in stimulated and stimulated, TTX-treated neurons. Stimulated sample: N = 75 synapses, TTX sample: N = 28 synapses. (**f**) Examples for SVs connected to fusion events. i: tomogram slice and zoom-in (lower panel), i’: isosurface of a fusion stalk with two additional SVs. Asterisks label fusion events, numbers display connected SVs, arrowheads display connecting filaments. Scale bars 20 nm. ii-iv: Exemplary tomogram slices of dilating (left panel) and collapsing (middle and right panels) fusion pores connected to SVs. Scale bars 20 nm. (**g**) Exemplary tomogram slices of stimulated synapses with prominent presynaptic filaments likely resembling actin. The measured distance between horizontal filament and active zone membrane was 17 nm (lower panel). Pink boxes indicate positions of zoom-ins with SVs connected to these filaments (right panels). Blue arrowheads indicate actin-resembling filaments, pink arrowheads indicate filaments connecting them to SVs. Scale bars left panels: 100 nm, right panels: 20 nm.

Particularly during fast and sustained neurotransmitter release, release sites need to be refilled with SVs. In our synapses, we observed structural features that may contribute to different modes of release site refilling. Firstly, we noticed that SVs were not only connected to each other via pleomorphic interconnectors, but also partially connected to fusion intermediates of different categories (**Fig. 5f**, also see^25,26^). Considering that filamentous connections between membrane-near SVs were shown to persist or even increase during SV fusion^25,26^, it is likely that also the linkers between SVs and fusion events are stable and/or strengthened during action potential-induced calcium influx. This way, fusing SVs may directly recruit new SVs for very fast release site refilling. However, we observed such connections only at a small fraction of fusion events. Secondly, we observed filamentous connections between SVs and cytoskeletal filaments like actin (**Fig. 5g**). In stimulated neurons with and without ongoing fusion, we occasionally observed horizontal, single stranded filaments in short distance (17-36 nm) to the cell membrane and preferentially at the borders of the AZ (**Fig. 5g, lower panel**). We further observed vertical and angular filaments spanning around and also within the cytosolic SV pool above the AZ (**Fig. 5g, upper and lower panel**), which were recently described as actin corral and actin rails, respectively^51^. Although we could not correlate the prevalence of these cytoskeletal filaments to the strength of the individual synapses, it is conceivable that actin spans around synaptic AZs to help organizing the enrichment and distribution of SVs for fast and sustained neurotransmitter release, as suggested recently^51,52^.

## Discussion

In summary, we have developed a workflow combining optogenetic stimulation of neurons with plunge freezing and cryo-ET to achieve a combination of highest possible temporal and structural resolution, paired with near-native cellular preservation and near-physiological stimulation. Although optogenetic plunge freezers have been developed before^53–55^, we could now verify the suitability of such a setup for *in situ* applications. In comparison to other approaches for time-resolved cryo-EM, typically involving the mixing and spraying of reactants onto EM grids^39^, we achieved a similar temporal resolution with optogenetics. The coupling of light pulse and cryofixation may thus not only be of interest for *in situ* experiments like ours but also for *in vitro* applications.

Our time-resolved cryo-ET workflow formed the basis for an in depth characterization of SV fusion *in situ*. While previous cryo-ET studies using chemical stimulation already showed examples of SV fusion intermediates in synaptosomes^25,26^, we were now able to categorize and quantitatively assess SV fusion states from fusion initiation to membrane integration. Overall, our observed fusion intermediates closely resembled fusion states recently described in all atomic MD simulations^11^ and will likely be of help for setting up future simulations and models. While there is a general consensus about the opening and collapsing of fusion pores, the mechanisms behind fusion initiation are still under debate and may vary between different cellular processes^3^. Our observations favor a model in which SV fusion in neuronal synapses is initiated by stalk formation, whereby the AZ membrane is only slightly invaginated when the spherical SV converts into a droplet shape. Although we and others^26^ observed AZ membrane invaginations below tethered SVs, we do not have experimental evidence that they directly lead to SV fusion. MD simulations described membrane invaginations preceding SV fusion beforehand and attributed them to membrane curvature-inducing functions of proteins such as synaptotagmin-1^56,57^. However, a direct role of synaptotagmin-1 in inducing membrane curvature has been questioned recently^10,12^. Our coarse-grained simulation likewise indicated that the induction of membrane curvature through proteins like synaptotagmin-1 may not be beneficial for fusion initiation. Instead, the observed slight membrane invaginations may originate from the zippering of the SNARE complex, as recently shown in an MD simulation of SNARE-mediated SV fusion without synaptotagmins^58^.

Beyond mediating membrane fusion, exocytic proteins were shown to be involved in the recruitment and the priming of SVs^6,40^. Overall, our findings support the idea that multiple tethers are preferentially formed between the AZ membrane and SVs in close membrane proximity^25–27^. Yet, we neither found differences in tether numbers between synapses with and without ongoing SV fusion nor between SVs with and without membrane invaginations. Considering that we observed fewer SVs in stimulated synapses primarily within a distance of 6 nm from the AZ membrane and membrane invaginations below SVs with an average distance of 6.4 nm, it is likely that most functionally primed SVs are also located here. Consequently, not the number of tethers but rather their molecular composition may correlate to the priming state of SVs^2^. Indeed, based on their size, not only assembled SNARE complexes together with synaptotagmin-1 and complexin, reflecting a protein interaction during priming, would fit into densities observed below membrane-near SVs^19,27,28^. At least in our example, Munc18 interacting with syntaxin-1, which reflects a pre-priming state, would fit equally well. Importantly, these observations need to be interpreted with caution: Due to the limited resolution of *in situ* studies like ours, the required structural resolution is missing to reliably fit the one or the other protein complex into the observed densities.

We further noticed that the number of membrane-proximal SVs in the subgroup of synapses without ongoing fusion was lower than expected. A possible explanation for this observation is that synapses with very fast responses (too fast to be captured with our setup) also showed higher fractions of MVR, leading to a stronger depletion of the membrane-proximal SV pool. Indeed, our observed probability for MVR was lower than described previously for comparable stimulation conditions^47^. However, also a transient dissociation of proximal SVs from the AZ membrane is conceivable, as previously suggested^13^.

## Material and methods

### Mass culture of mouse primary hippocampal neurons

#### Astrocyte feeder culture

All experimental procedures involving the use of mice were approved by the Animal Welfare Committee of the Charité-Universitätsmedizin Berlin and the Berlin State Government. Astrocytic and neuronal mass cultures were prepared from P0-P2 C57/BL6/N mice of either sex. To prepare astrocyte feeder layers, mice were decapitated and cortices were isolated in cold HBSS-HEPES. The tissue was digested in 0,05% trypsin-EDTA for 15-20 min at 37°C followed by manual trituration. The isolated astrocytes were transferred to T75 flasks and cultured for two weeks in Dulbecco’s modified Eagle medium supplemented with 10% fetal calf serum (FCS), 10,000 U/ml penicillin and 10,000 µg/ml streptomycin (DMEM) at 37°C and 5% CO2. Confluent astrocytes were trypsinated (0,05% trypsin-EDTA) and seeded on collagen/poly-D-lysine coated coverslips (for RT experiments) or coated wells (for cryo experiments and banker cultures) in a density of 75,000 cells/well (six-well plates). The astrocytes were cultured for an additional week before FUDR (8.1 mM 5-fuoro-2-deoxyuridine and 20.4 mM uridine in DMEM) was added to arrest glia proliferation.

#### Neuronal culture on coverslips

For live imaging, RT confocal imaging and electrophysiological recordings, primary hippocampal neurons were used as co-cultures with astrocytes or separated from astrocyte feeder layers as banker cultures. Since we did not find differences in the responsiveness of neurons in the two different culture systems, we pooled data from mass cultures and banker cultures. To isolate neurons, hippocampi were dissected in cold HBSS-HEPES and digested using 20 U/ml papain for 45 min at 37°C, followed by manual trituration. For co-cultures, isolated neurons were seeded directly on 1-2 weeks old astrocyte feeder layers in a density of 3·10^4^ - 5·10^4^ neurons/well. For banker cultures, neurons were seeded on coverslips coated with collagen/poly-D-lysine and ornithine (5·10^4^ neurons/well), and the coverslips were transferred to well plates containing astrocyte feeder layers after neurons were allowed to adhere to the coverslips for 1 h. Neurons were cultured for 14-18 days at 37°C in neurobasal-A medium containing 2% B-27, 1% Glutamax, 10^5^ U/ml penicillin and 10^5^ µg/ml streptomycin (NBA), lentiviruses and AAVs were added on DIV2-3.

#### Neuronal culture on EM grids

For culturing neurons on EM grids, mesh pedestals were 3D printed and placed on top of astrocyte feeder layers. DMEM medium was replaced by NBA medium. Quantifoil R3.5/1 AU holey carbon grids (400 mesh, 200 mesh and 200 mesh finder grids) were cleaned with chloroform and acetone, followed by glow-discharging. Directly afterwards, the grids were placed on droplets of collagen/poly-D-lysine coating solution and incubated for 20 min under UV light and 1-2 hours at 37°C. The grids were washed in PBS overnight and then transferred to the well plates containing the astrocyte feeder layers and pedestals, 3 grids per well. Primary hippocampal neurons were seeded in a density of 1.5·10^5^ - 2·10^5^ cells/well, viruses were applied on DIV2-3.

### Live imaging of biosensors

Live imaging of neurons expressing the glutamate sensors iGluSnFR, iGluSnFR3-PDGFR, or iGluSnFR3-GPI was performed at elevated temperature (∼32-34°C) using an inverted microscope (Olympus IX51) with a custom-built in-line heating system and a 60x water immersion objective with a heated-collar (Warner Instruments). Cells were perfused with high-calcium extracellular solution containing the following: 140 mM NaCl, 2.4 mM KCl, 10 mM HEPES, 10 mM glucose, 4 mM CaCl_2_, and 1 mM MgCl_2_ (300 mOsm; pH 7.4). 6 µM NBQX, 30 µM bicuculline, and 10 μM AP-5 were added to block neuronal network activity of the mass culture. Action potentials (2 ms depolarization) were induced using a field stimulation chamber (Warner Instruments), Multiclamp 700B amplifier, and an Axon Digidata 1550B digitizer controlled by Clampex 10 software (all Molecular Devices). To assess the suitability of the biosensors for our cryo-ET workflow, two action potentials with an inter-stimulus interval of 25 ms were induced, which corresponds to the minimal time between first and second stimulus of the optogenetic plunge freezer (limitation due to the bottom of the incubation chamber of the Vitrobot). Samples were illuminated by a 490 nm LED system (CoolLED) with an exposure time of 10 ms, images were captured with an andor iXon Life 897 camera (Oxford instruments) at a frame rate of 40 fps.

### Optogenetic stimulation and vitrification

For optogenetic stimulation of neurons, a plunge freezer (Vitrobot Mark IV, Thermo Fisher Scientific) was equipped with a high intensity LED (Schott LLS3, wavelength 470 nm), from which one PMMA optical fiber with PVC insulation (inner diameter 3 mm) reached inside and one below the chamber of the Vitrobot. The end points of the optical fibers were installed with a distance of max. 5 mm to the path of the plunge frozen EM grids and illuminated the grids entirely. The LED was controlled by an infrared sensor that detected the downward motion of the tweezer holding the EM grid. The grid was illuminated twice for 5 ms: the first light pulse was applied within the chamber, max. 100 ms before freezing, and the second light pulse was started approximately 7 ms before the sample reached the liquid ethane. The correct timing and illumination were confirmed using the super-slow-motion mode of a Samsung Galaxy S20 camera with a frame rate of 960 fps.

At DIV16-18, each EM grid containing neurons was briefly washed in a high-calcium solution containing 140 mM NaCl, 2.4 mM KCl, 10 mM HEPES, 10 mM glucose, 4 mM CaCl_2_, 1 mM MgCl_2_, 3 μM NBQX, and 30 μM bicuculline (∼300mOsm; pH7.4) pre-warmed to 37°C and directly transferred to the plunge freezer. 4 µl of high-calcium solution supplemented with 10 nm BSA-gold (Aurion, OD∼2) were applied on the grid prior to blotting for 12-16 s (backside blotting, blot force 10) at 37°C and a relative humidity of 80%. The grids were plunge-frozen in liquid ethane and stored in liquid nitrogen until further use. For TTX-treatment, grids were incubated in high-calcium solution containing 1 µM TTX for 1-2 min prior freezing.

### Cryo-confocal microscopy

#### Data acquisition

After plunge freezing, EM grids were clipped into autogrids and transferred to a TCS SP8 cryo-confocal microscope equipped with a 50x CLEM cryo-objective, NA 0.9 (Leica Microsystems). Overviews of each grid were acquired in brightfield and fluorescence mode; the reflective mode was used on a subset of grids to estimate the ice thickness. The iGluSnFR fluorescence signal was used to confirm overall successful grid stimulation. For the comparison of iGluSnFR fluorescence intensities without and with stimulation, only the fluorescence signal of the fluorophore attached to ChR2 (YFP or mScarlet) was used to select regions of interest for subsequent cryo-confocal microscopy to avoid bias. Cryo-confocal stacks (z-steps 0.5 µm, pixel size 0.11 µm) were acquired with optimized filter settings of the HyD detector to minimize YFP or mScarlet crosstalk with the GFP signal. From grids intended for correlative confocal and electron microscopy, only few confocal stacks were acquired and the overall acquisition time per grid was limited to 30 min to avoid strong ice contaminations.

#### Analysis of cryo-confocal microscopy and correlation with cryo-electron tomography

Fluorescence intensities in cryo-confocal stacks of unstimulated, stimulated, and TTX-treated plunge-frozen neurons were compared using fiji software^59^. Maximum intensity z-projections of each confocal stack were generated and 300x300 pixel regions of interest (ROIs) containing individual neurites were extracted. The mean fluorescence intensity (**Fig. 2d**) was measured per ROI and averaged per confocal stack. To compare the distribution of fluorescence intensities (**Fig. 2c**), fluorescence intensity histograms were generated for each ROI. A threshold of 15 was applied for background subtraction and the pixel counts per intensity were normalized to the total pixel number per ROI. These relative intensity histograms of ROIs were averaged per confocal stack. The threshold of 70 for high-intensity pixels was visually assessed in stimulated samples. The fraction of pixels >70 (**Fig. 2e**) was calculated per ROI and averaged per confocal stack. The correlation of fluorescence and TEM (**Fig. 2f**) was performed manually using the navigator of the Leica lasx software and fiji.

### Cryo-electron tomography

#### Data acquisition

Cryo-ET data collection of optogenetically stimulated neurons cultured on EM grids was performed on a Titan Krios G3i electron microscope (Thermo Fisher Scientific) equipped with a K3 direct electron detector with BioQuantum energy filter (Gatan) and operated at 300 kV. Tilt series were typically acquired with 10 frames per tilt at a magnification of 15,000x and a pixel size of 3.2 Å in superresolution mode using PACE-tomo^60^. Tilt angles ranged from −50° to +50° and 2° angular increment in a dose-symmetric^61^ tilt-scheme. The defocus values ranged from −3 to −6 µm and the total electron dose was 106-125 e^-^/ Å².

#### Tomogram reconstruction

Tomograms used for analysis were reconstructed semi-automatically using the tomoBEAR^62^ pipeline: Aligned frames were motion-corrected using MotionCor2^63^. The tilt series alignment was performed by DynamoTSA^44^ and manually refined using 10 nm gold fiducial markers in IMOD^64,65^. For each projection, defocus values were measured by Gctf^66^, and CTF correction was performed using the IMOD command ctfphaseflip^67^. Four-times binned 3D reconstructions (final pixel size 12.28 Å) from CTF-corrected, aligned stacks were obtained by weighted back projection in IMOD.

In total, 312 tilt series of stimulated neuronal samples and 95 tilt series of TTX-treated neuronal samples were reconstructed with tomoBEAR and manually screened for synapses, which were only recognizable after reconstruction. We visually identified synapses as presynaptic boutons filled with SVs and a synaptic AZ, a synaptic cleft of 10-30 nm width, and a postsynaptic bouton with visible postsynaptic density. Based on these criteria, we identified 75 synapses in the stimulated samples and 28 synapses in the TTX-treated sample that were used for further analysis. For cryo-ET analyses of stimulated synapses, we did not analyze each freezing/grid individually but pooled synapses of three grids/two freezings because we did not note any significant differences in numbers of SVs or putative fusion events between them.

### Segmentation and analysis of cryo-electron tomography data

#### Segmentation

Automated segmentations of synapses (**Fig. 1** and **2g**) were performed with MemBrain v2^68^: Tomograms were denoised and corrected for the “missing wedge” effect using IsoNet^69^, 4- times binned and lowpass filtered using IMOD. Membranes (intracellular organelles and plasma membranes) were segmented automatically with MemBrain v2 and corrected manually using Amira (Thermo Fisher). The segmentations were re-colored and aligned with tomogram slices using ChimeraX. Manual segmentations of AZs (**Fig. 4a**) were made with IMOD using 4-times binned, IsoNet-corrected tomograms.

#### Morphometric characterization of membrane rearrangements

All synapses were screened for membrane rearrangements potentially resembling fusion events. From these ROIs, 4-times binned subtomograms with a box size of 200x200x200 pixels (pixel size 12.28 Å) were generated using *dynamo_catalogue*^70^. As a quality control and to avoid bias, the original tomograms were screened and ROIs were preselected by the first author. The second author double-checked all positions independently and generated the subtomograms. Initial ROIs were excluded if the putative fusion event was not located at an AZ with recognizable postsynaptic density, if a halo resembling a clathrin coat around the pore was visible, or if the resolution of the tomogram was poor (see **Suppl. Fig. 6** for examples of excluded ROIs). The subtomograms were denoised and corrected for missing wedge effects using IsoNet. These denoised subtomograms were used for the classification of putative fusion states. In addition, we double-checked the correct classification for a subset of ROIs using the original (un-denoised) dataset. Based on our observations, we defined 7 categories of membrane rearrangements at the AZ (opposing the postsynaptic density) and two additional states of SVs in membrane proximity (see **Suppl. Table 1** for morphological characteristics of each category).

Center slices of putative fusion events were exported from IMOD as tiff files. Size measurements (**Fig. 3a**) were performed in fiji: the horizontal diameter of tethered SVs with membrane invagination, stalk formation, closed, and open fusion pores was measured at the widest region of the respective SV/fusion event between the outer borders of the lipid bilayers. The width of dilating and collapsing fusion pores, as well as bumps was measured between the two positions of the AZ membrane where inward curvature was observed. For the distance of tethered SVs and SVs during stalk formation, the space between lipid bilayers of SV and membrane was measured, whereby membrane in- and evaginations were interpolated. For the height of stalks, closed, open, dilated, and collapsing fusion pores, the distance between AZ membrane outer (upper) border and the outer border of the fusion event membrane at the highest position was measured, whereby invaginations of the AZ membrane (stalk formation, closed fusion pore) were interpolated. The height of the neck of the closed fusion pore was defined as the region in which both walls of the pore (in 2D) were in direct contact. The pore width of the open fusion pore was defined as the space between lipid bilayers of the pore walls at the narrowest position of the neck. The thickness of membranes at collapsing fusion pores was measured at the top of the pore and next to the pore base.

#### Quantification of fusion events, multivesicular release and bumps

Based on our definition of membrane rearrangements, we quantified numbers of observations per category. We first quantified synapses containing at least one of these events (**Fig. 3d**). If synapses contained more than one event, only the event closest to category 4 (as center) was used to define the overall state. Since stimulated synapses contained more events of categories 2-6 than TTX-treated synapses, we defined these states as “ongoing SV fusion”. Based on this definition, we had three groups of synapses: synapses with ongoing fusion, synapses with bump(s), and synapses without membrane rearrangements (**Fig. 3d-e**). Synapses containing at least two fusion events (categories 2-6) were counted as MVR synapses (**Fig. 5c,d**). Additionally, we counted each fusion event individually (**Fig. 3f**).

#### Quantification of SV distances and tethers

The quantification of distances and filamentous tethers connecting SVs and the AZ membrane (**Fig. 4d-4g**) was performed in IMOD using IsoNet-corrected tomograms. To make sure that the observed filaments were not a denoising artifact, we double-checked our annotations in a subset of ROIs using the original (un-denoised) dataset. For distance measurements, we first labeled all membrane-near SVs of each synapse above the AZ up to a distance of 24 nm at lower zoom and then measured their exact distances at higher zoom using the IMOD measuring tool. The distance between SV and AZ membrane was measured at the center slice of the SV and defined as the space between lipid bilayers of SV and membrane. We analyzed distributions of membrane-near SVs per synapse, whereby synapses containing no or only one SV were excluded. For this, we binned distances in 2 nm steps. Based on this distribution, previous reports^13,50^ and our observation that membrane invaginations were visible below SVs with an average distance of 6.4 nm, we further defined three subpools of SVs: membrane-proximal SVs with a max. distance of 6 nm, intermediate SVs with a distance of 6-12 nm, and distal SVs with a distance of 12-24 nm.

Only synapses with very good structural resolution (high signal-to-noise ratio, clearly visible membrane bilayers, etc.) were used for tether analysis. Tethers were defined as vertical or angular filamentous connections between SV and AZ membrane. We quantified tethers per SV manually, whereby we went back and forth in z direction several times and at different zooms. The slicer window was additionally used to rotate SVs around the x and y axis. Only if a filament was visible on multiple z slices, it was counted.

### Simulation

The coarse-grained model of the SV and the AZ includes particle-based representations of bilayer membranes, curvature-inducing proteins, and bead and spring models of tether and SNARE proteins. For the membranes, we used the membrane model developed by Sadeghi and Noé^71^. This two-particle-per-thickness coarse-grained model is parameterized to mimic the mechanics of a fluid membrane with specified bending rigidity, has tunable in-plane viscosity, and is coupled with a hydrodynamics model that reproduces the out-of-plane kinetics of membranes in contact with solvents of prescribed viscosity^72,73^.

To parameterize the membrane model, we used reported values of the elastic response of SVs to indentation forces in atomic force microscopy (AFM) measurements^74^. We used these values in conjunction with a theoretical model, initially developed for the AFM indentation of influenza virus envelopes^75^, that relates the overall stiffness (or spring constant) of a spherical vesicle to the bending rigidity of its membrane. We found a mean membrane bending rigidity of 0.8⨯10^-19^ J, which is well within range of values obtained for lipid bilayers^76^.

We modeled the SV in the initial state as a sphere with the outer diameter of 42.5 nm, to reflect the mean values obtained from tomograms (**Fig. 3a**), and added a harmonic volume-preserving potential to its outer leaflet particles. This potential acts against any changes in the enclosed volume, while allowing for otherwise arbitrary deformations. The plasma membrane underneath the vesicle is modeled as a planar square membrane patch of 180 nm in side length. The simulation box is coupled in-plane to a stochastic barostat that controls the lateral pressure components around zero.

We modeled the curvature-inducing proteins in the AZ (**Fig. 3g**) via a force field masking mechanism that allows for tagged particles to locally modify the interparticle interactions using Monte Carlo moves, while letting these particles freely diffuse within the membrane. The modified force field reflects a preferred signed curvature (upward/downward) around these particles^77^. We used reported values of membrane curvatures upon binding cyclic peptides derived from synaptotagmin-1 C2B domain to assign preferred local curvatures to these particles^78^.

The tether proteins (**Fig. 3g**) are included as fixed-length elements initially formed between particles on the vesicle to anchor particles on the plasma membrane. The position of these tethers are decided randomly at the initial state. We incorporated a harmonic angle-bending potential that, when activated, exerts a torque to rotate the tethers about their anchor point on the plasma membrane, in effect pulling the SV toward the AZ.

We included chain-like representations of two sets of SNARE proteins, namely v-SNARE on the SV and t-SNARES in the AZ. The sizes of the chains roughly match the overall structure of SNARE proteins synaptobrevin, syntaxin, and SNAP-25 in the zippered complex. We included selective short-range attractive pairwise interactions between beads that form v-SNARE and t-SNARE chains such that they prefer to match one-to-one in the correct zippered configuration. Spatial exclusion, modeled via soft harmonic repulsions, energetically prohibits other conformations. We chose the copy number of SNARE proteins based on the reported proteomics data for SVs^79^, and calibrated the strength of attractive interactions to match the data on zipping/unzipping forces measured with magnetic tweezers^80^.

To test the effect of curvature-inducing proteins in the AZ, we developed models with 0, 10, 20, and 30 copies of these proteins. For each model, we started 5 simulation replicas, each with different random distributions of tether, SNARE, and membrane-curving proteins. We used anisotropic Brownian dynamics with a timestep of 0.1 ns to simulate the motion of all the particles in the system, and assigned a cytosolic viscosity of 2.21 cP^81^ to calculate particle mobilities in our hydrodynamic coupling method^73^.

At the start of each simulation, the system is allowed to relax for 5 μs, with the torque on the anchor points of tether proteins disabled, effectively having the SV floating at constant distance with the AZ (**Suppl. Fig. 8a**). Afterwards, the tethers are activated, pulling the vesicle toward the AZ. The gap between the two membranes in their closest approach is continuously monitored. When the gap falls below 6 nm, we initiate the force field masking mechanism for curvature-inducing proteins, which results in curvature being developed in the AZ membrane. Each simulation thus continues for another 100 μs to follow the docking dynamics (**Suppl. Fig. 8**).

Trajectories are obtained by sampling the positions of all the particles at 100 ns intervals. All the subsequent trajectory post-processing, data analysis, and plotting is done through Python scripts, using Numpy^82^ and Matplotlib^83^ software packages. The 3D visualization is done via the software package Visual Molecular Dynamics (VMD)^84^.

### Statistics and data representation

Fluorescence microscopy data were analyzed using fiji (fluorescence intensity means and histograms) and python, cryo-ET data were quantified using IMOD and fiji. Graphpad prism was used for statistical tests. P-values were defined as follows: * p<0.05, ** p<0.01, *** p<0.005. Graphs were generated using the python packages Seaborn^85^ and Matplotlib^83^ and optimized for visualization using Affinity Designer 2. Isosurfaces of bandpass-filtered tomograms and segmentations were visualized using ChimeraX, whereby the “hide dust” function was used. Only for subtomogram averages, the erase tool was used to remove artifacts that were not in direct contact with the particle and introduced/ amplified by C61 symmetry. Density maps of atomic models with a resolution of 10 Å were generated with the ChimeraX function molmap and manually fitted into cryo-ET densities. If not stated differently, data are represented as mean ± standard error of the mean (sem). In violin plots and bar graphs, the center line depicts the median, the upper and lower lines/borders of the box display the 25% and 75% percentiles. Whiskers in box plots indicate the 10-90% percentile range in all graphs except Suppl. Fig. 8 (here min. to max.). XY plots show lines connecting means and the semitransparent areas indicate the sem.

## Contributions

JK and CR designed the study, CR supervised the project. JK and MaS conceived the setup for optogenetic plunge freezing. CAD designed and built the optogenetic freezing device. JK developed a protocol to culture neurons on EM grids. JK and MaS performed plunge freezings. JK and TS acquired cryo-ET data. UK processed cryo-ET data. JK and UK analyzed cryo-ET data and segmented tomograms. MoS generated the computational model, performed the corresponding simulations and analyses. JK acquired and analyzed cryo-confocal microscopy and CLEM data. LI and JK acquired live imaging and RT confocal microscopy data. JK and LI analyzed live imaging data. ML acquired and analyzed electrophysiology data. JK designed figures and prepared the manuscript. All authors reviewed the manuscript.

## Acknowledgements

We thank Heike Lerch, Berit Söhl-Kielczynski and members of the Rosenmund lab for technical assistance. We thank Marion Weber-Boyvat, Pascal Fenske, Melissa Herman and Severin Dicks for help with live imaging and the respective data analysis. We thank Metaxia Stavroulaki for help with cryo-confocal microscopy and CLEM and Timo Flügel and Simon Lauer for help with plunge freezing. We thank Artsemi Yushkevich for help with data processing. We thank Melissa Herman for critical reading of the manuscript. We thank the Viral Core Facility of the Charité-Universitätsmedizin Berlin for lentivirus and AAV production.

