## Supplementary figures, tables and methods for "Dynamic nanoscale architecture of synaptic vesicle fusion in mouse hippocampal neurons"

#### Supplement

Suppl. Fig. 1: Assessment of action potential induction efficacies of three Channelrhodopsin-2 variants.

Suppl. Fig. 2: Banker culture of mouse hippocampal neurons on EM grids.

Suppl. Fig. 3: Setup and timing of optogenetic plunge freezer.

Suppl. Fig. 4: Live-imaging of fluorescent activity sensors.

Suppl. Fig. 5: Correlation of iGluSnFR3 cryo-confocal microscopy and EM of optogenetically stimulated neurons.

Suppl. Fig. 6: Examples of putative fusion events that were excluded.

Suppl. Fig. 7: Ultrastructural characteristics of putative SV fusion intermediates.

Suppl. Fig. 8: Coarse-grained simulation of SV approximation.

Suppl. Fig. 9: Distributions and tethering of SVs in stimulated and control synapses.

Suppl. Fig. 10: Exemplary alignment of vesicular tether sizes and exocytic proteins.

Suppl. Table 1: Morphological characteristics of SVs and fusion states.

Suppl. Table 2: Viral constructs.

Supplementary methods

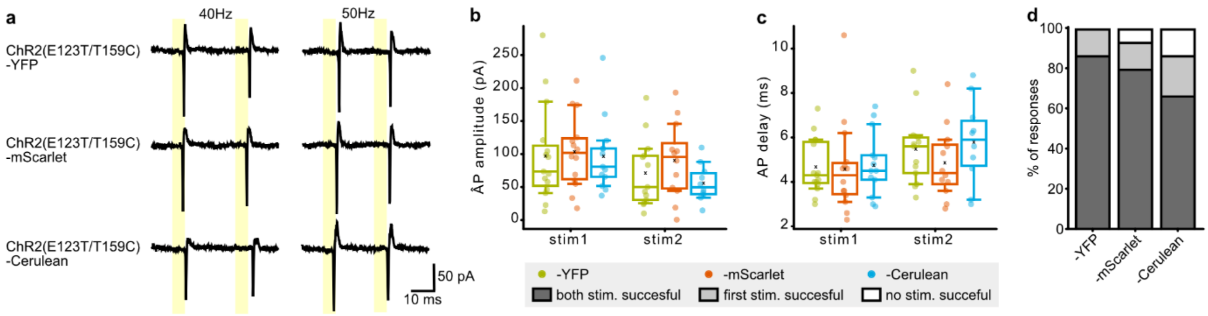


**Suppl. Fig. 1: Assessment of action potential induction efficacies of three Channelrhodopsin-2 variants.** (**a**) Exemplary light-induced responses to 40 Hz and 50 Hz paired pulses of ChR2 variants measured in cell attached, patch clamp recording mode, light pulse duration 5 ms. (**b**) Action potential amplitudes in response to first and second light pulse at 40 Hz stimulation. (**c**) Delay between light onset and action potential induction for first and second light pulse at 40 Hz stimulation. (**d**) Success rate at 40 Hz stimulation. Dark gray: both stimuli successful, light gray: only first stimulus successful, white: stimulation not successful. ChR2(E123T/T159C)-YFP N = 15 cells, ChR2(E123T/T159C)-mScarlet N = 14 cells, ChR2(E123T/T159C)-Cerulean N = 13 cells.

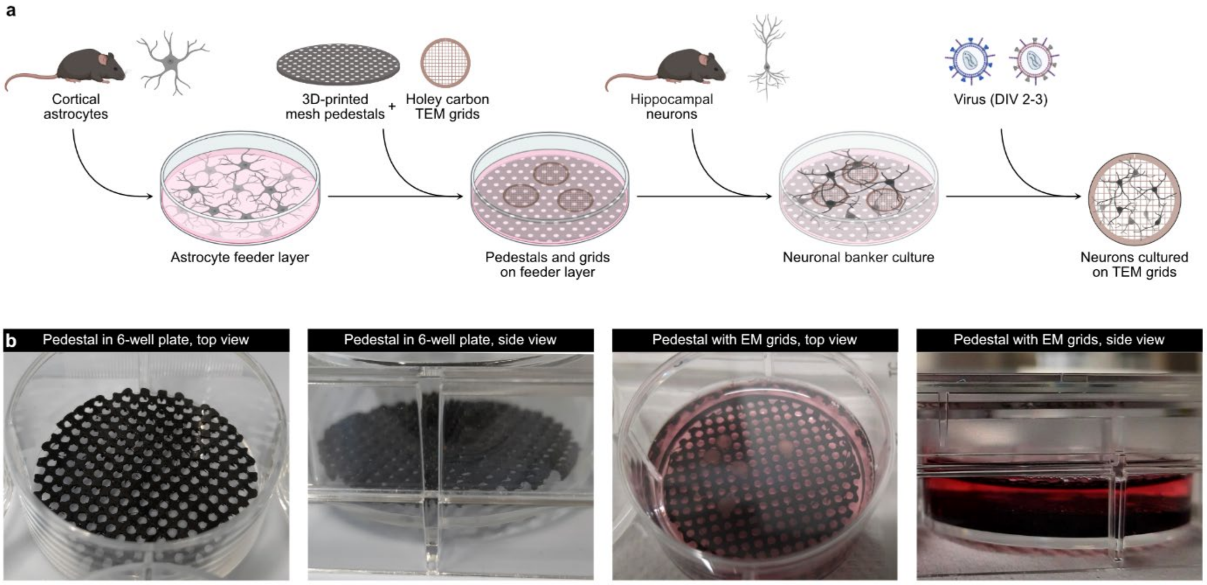


**Suppl. Fig. 2: Banker culture of mouse hippocampal neurons on EM grids.** (**a**) Schematic illustration showing the individual steps to culture neurons on EM grids. A feeder layer of mouse cortical astrocytes is seeded on 6-well plates and cultured for one week. 3D-printed mesh pedestals and glow-discharged, coated holey carbon grids are placed on top of the astrocyte feeder layer. Mouse hippocampal neurons are seeded and cultured for additional two weeks, viral constructs are added on neuronal day in vitro (DIV) 2-3. (**b**) Images of 3D-printed mesh pedestals in a well of a 6-well plate without (left) or with (right) cell culture medium and EM grids. The schematic illustration in (a) was generated with BioRender.

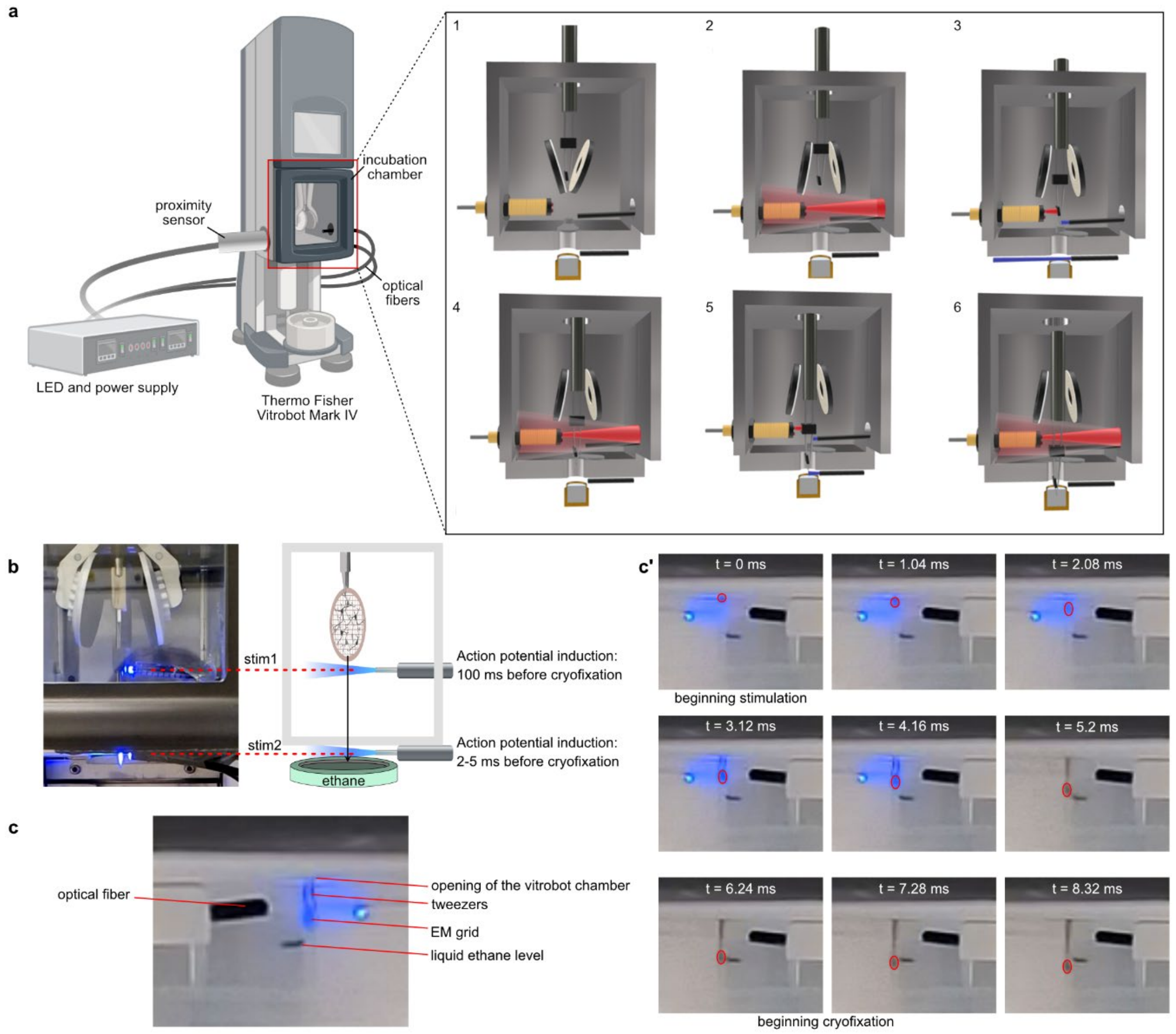


**Suppl. Fig. 3: Setup and timing of optogenetic plunge freezer.** (**a**) Schematic illustration of a Vitrobot Mark IV plunge freezer equipped with an LED and two optical fibers reaching inside and below the incubation chamber. The on-time of the LED is controlled with a proximity sensor. During grid loading and blotting, the LED is switched off (panel 1). As soon as the proximity sensor is activated by markers attached to the tweezers, the LED is switched on and blue light illuminates the path of the plunging EM grid (panel 2). The first light pulse is applied inside the chamber (panel 3), the second light pulse is applied below the chamber (panel 5). Between the two pulses (panel 4) and after the second pulse (panel 6), the LED is switched off. (**b**) Image of the Vitrobot chamber with two optical fibers when the LED is switched on. The timing between first light pulse and cryofixation in liquid ethane can be varied depending on the position of the optical fiber. For our experiments, we selected a time of ~ 100 ms between light pulse and freezing. (**c** and **c'**) Images of high-speed camera (frame rate 960 fps) capturing the illumination of an EM grid and the subsequent cryofixation, which is indicated by a black horizontal line at the level of liquid ethane (c). The illumination for optogenetic stimulation starts 6-8 ms before cryofixation. Taking into account that the delay between light onset and action potential induction was 3.9 - 5.9 ms for ChR2(E123T/T159C)-YFP, most action potentials were induced approximately 2-5 ms before cryofixation (c').

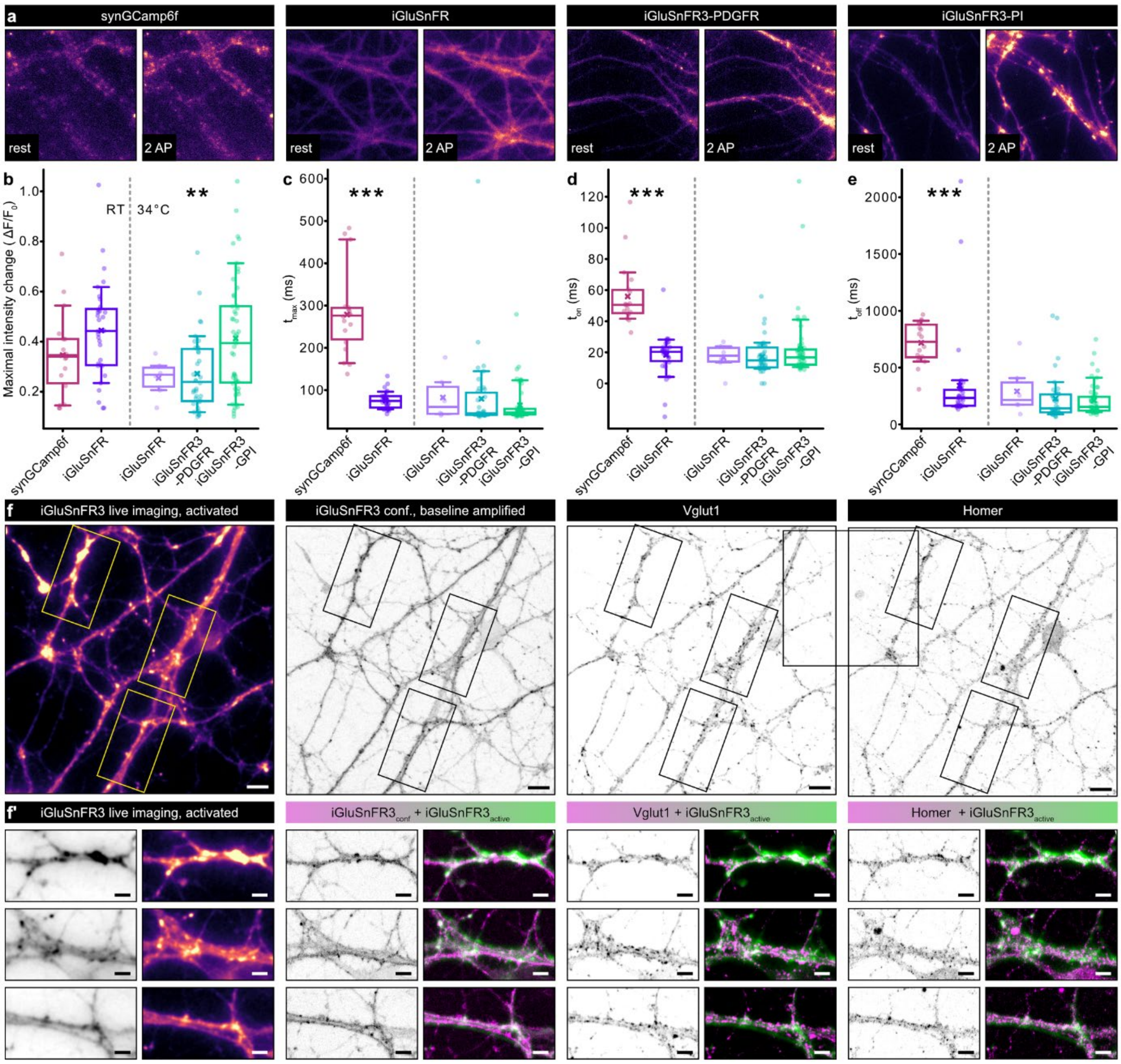


**Suppl. Fig. 4: Live-imaging of fluorescent activity sensors.** (**a**) Exemplary micrographs showing the activity sensors synGcamp6f, iGluSnFR, iGluSnFR3-PDGFR, and iGluSnFR3-GPI at rest and after application of one (synGcamp6f) action potential (AP) or two APs with a frequency of 40 Hz. (**b**) Maximal fluorescence intensity changes of the four activity sensors. (**c**) Delay between first action potential and time point with maximal fluorescence intensity. (**d**) Rise time until 50% of the maximal fluorescence intensity are reached. (**e**) Decay time until the fluorescence intensity decreased to 50% of the maximal intensity. (b-e) synGCamp6f and iGluSnFR (left boxplot) were recorded at room temperature, iGluSnFR (right boxplot) and both iGluSnFR3 constructs were recorded at a near-physiological temperature of 34°C. Box plots indicate mean, 25% and 75% quartiles, whiskers reach from 10-90% of data points. The x represents the mean. ** $p < 0.01$, *** $p < 0.001$. (**f**) Correlation of iGluSnFR3-GPI live imaging and *post hoc* immunostaining using antibodies against Vglut1 and Homer. Scale bars 10 µm. (**f'**) Zoom-ins to individual neurites and overlay of the iGluSnFR3 live imaging signal with iGluSnFR3 baseline (left), Vglut1 (middle) and Homer (right). Scale bars 5 µm.

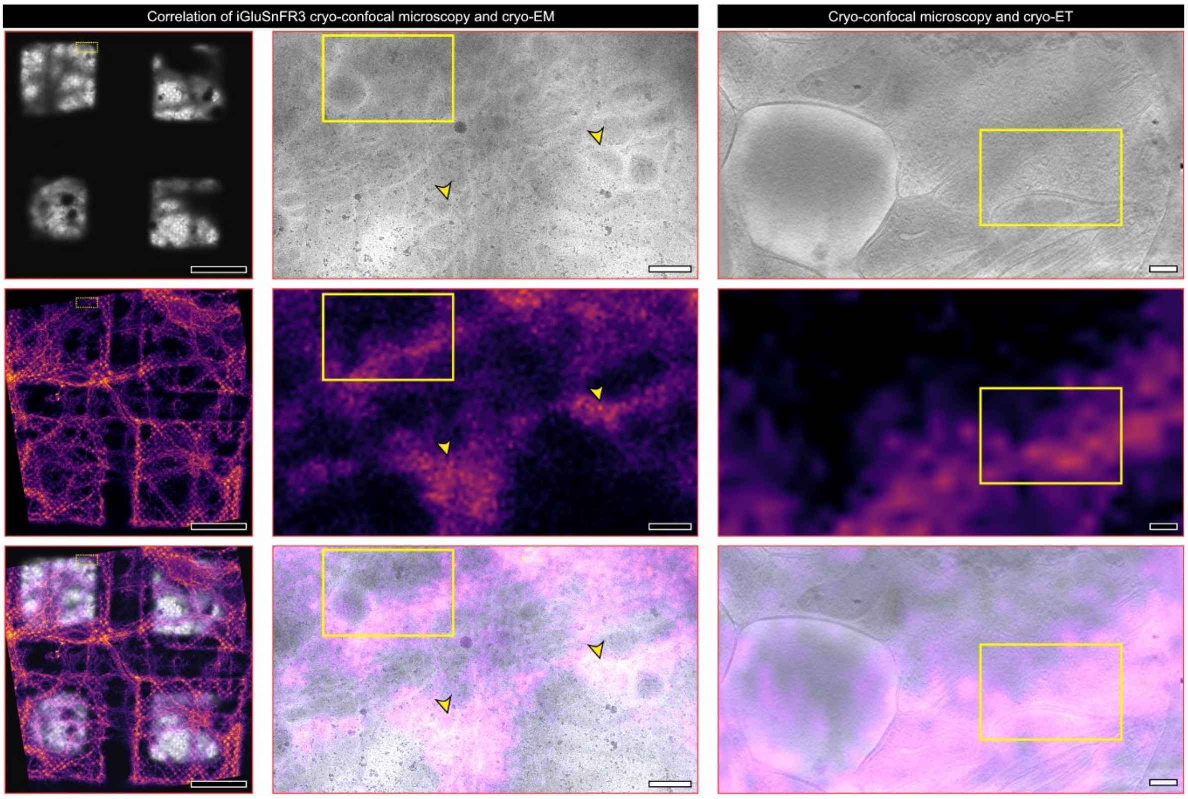


**Suppl. Fig. 5: Correlation of iGluSnFR3 cryo-confocal microscopy and EM of optogenetically stimulated neurons.** The top row contains TEM images (left, middle column) or tomogram slices (right column), yellow arrowheads and boxes point to regions with a high abundance of putative synaptic boutons. The middle row contains maximum intensity z-projections of cryo-confocal micrographs of the glutamate sensor iGluSnFR3. Here, the arrowheads and boxes point to areas with comparatively high fluorescence intensity, which indicates glutamate release. The bottom row contains overlays of fluorescence and EM signals. Scale bars left: 50 µm, middle: 500 nm, right: 200 nm.

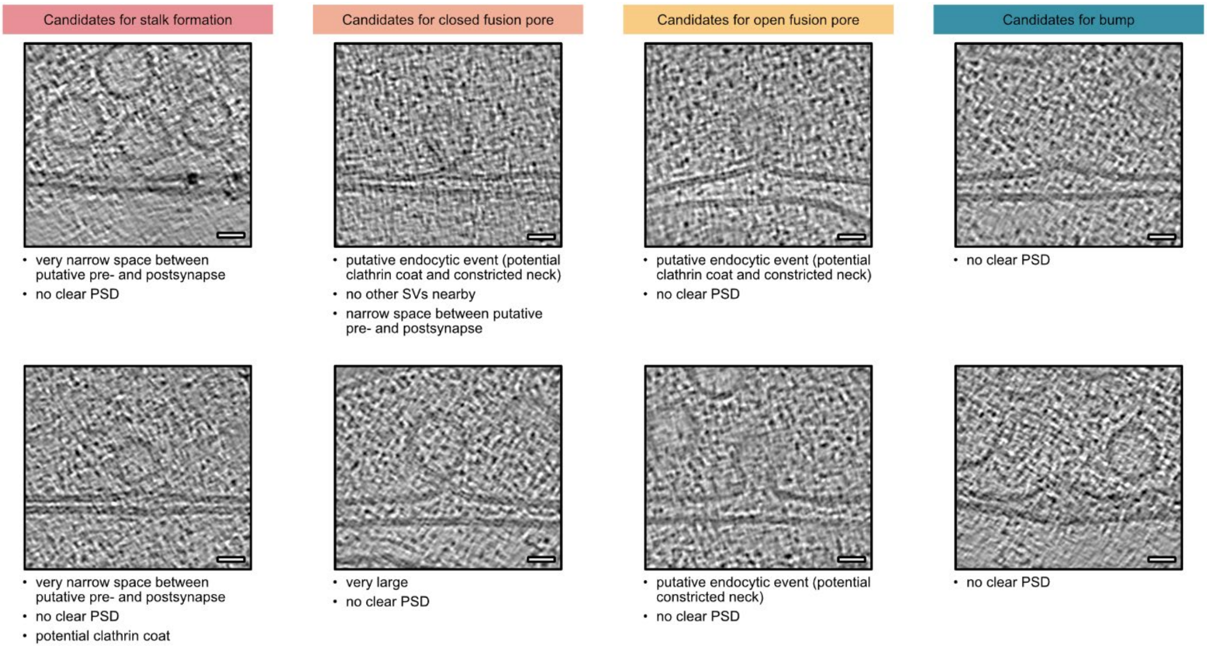


**Suppl. Fig. 6: Examples of putative fusion events that were excluded.** Tomogram slices of candidates for stalk formation, closed and open fusion pores and bumps. These and similar putative fusion events were excluded from the analysis because no postsynaptic density was visible, because the synaptic cleft was very narrow, or because a halo resembling a clathrin coat was visible. Scale bars 20 nm.

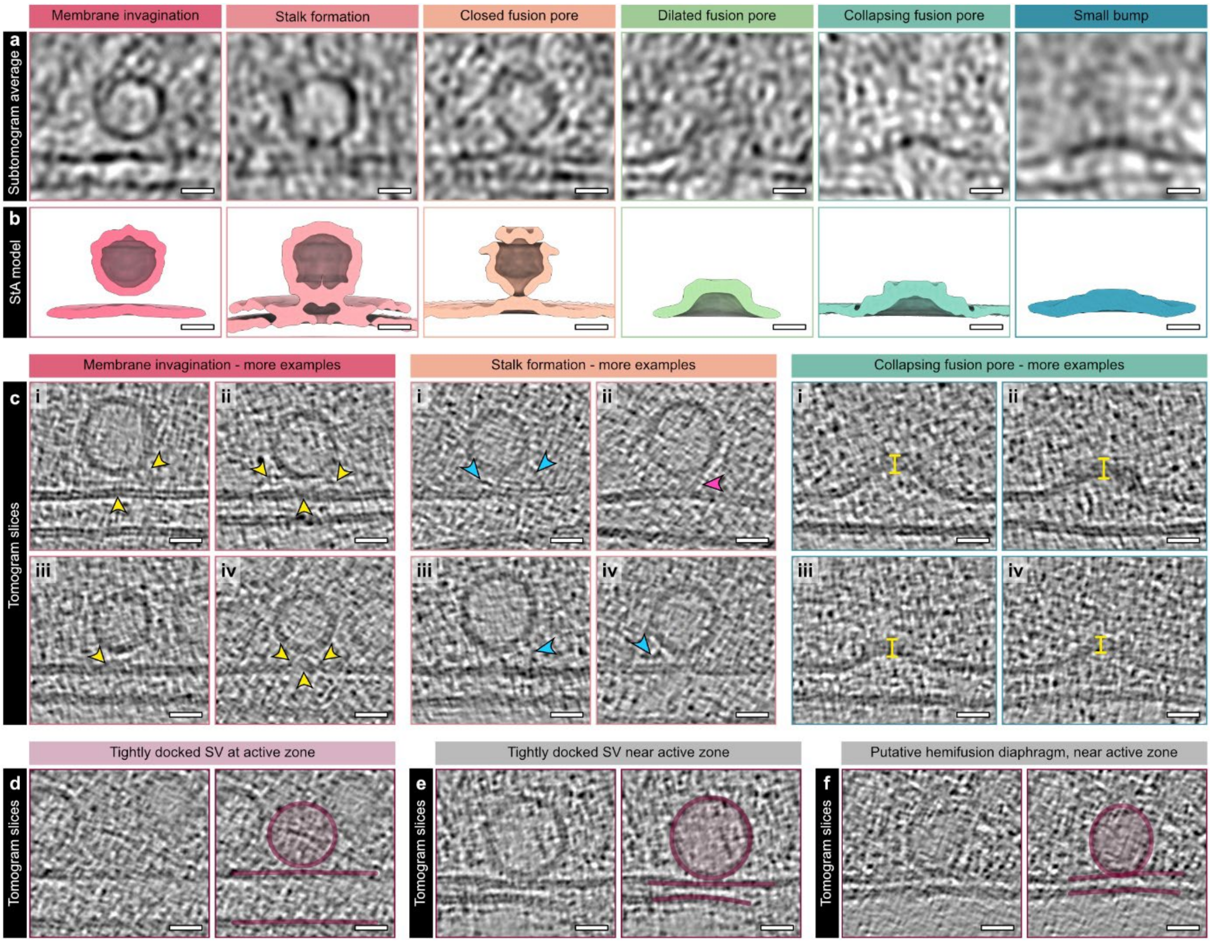


**Suppl. Fig. 7: Ultrastructural characteristics of putative SV fusion intermediates.** (**a**) Subtomogram averaging of putative membrane rearrangements during SV fusion. Membrane invagination (state 1) N = 5 averaged subtomograms, stalk formation (state 2) N = 8, closed fusion pore (state 3) N = 4, dilating fusion pore (state 5) N = 5, collapsing fusion pore (state 6) N = 6, small bump (state 7) N = 10 averaged subtomograms. (**b**) C61-symmetrized isosurfaces of the subtomogram averages of panel a. (**c**) Additional examples of SVs with membrane invaginations, stalk formations, and collapsing fusion pores. Not only below tethered SVs but also during stalk formation, only slight invaginations of the active zone membrane are visible (yellow arrowheads). During stalk formation, the space between SV and AZ membrane appeared partially blurry (pink arrowhead in example ii). Filaments were not only observed as tethers between the SV and the AZ membrane, but also attached to fusion events and reaching into the cytosol, as indicated by blue arrowheads. The top membrane of collapsing fusion pores was thicker than the AZ membrane (yellow lines). (**d**) Within all tomograms, only one example of a tightly docked SV at an AZ membrane was observed. (**e**) Additionally, a second tightly docked SV and a putative hemifusion diaphragm (**f**) were observed at membranes that were not opposed to a postsynaptic density, meaning not at an active zone. All scale bars 20 nm.

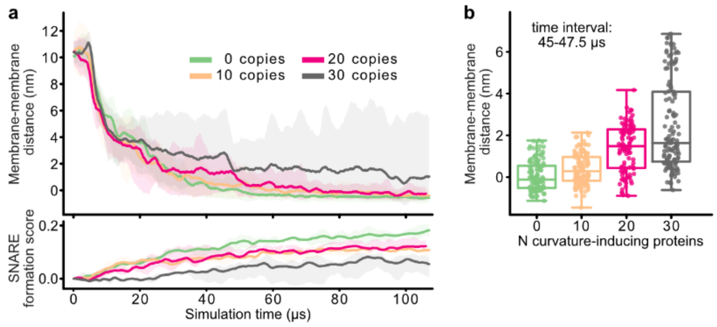


**Suppl. Fig. 8: Coarse-grained simulation of SV approximation.** (**a**) Top: vertical distance between the SV and AZ membranes, measured throughout the simulation, for simulations with the given copy numbers of curvature-inducing proteins in the AZ. Distance values are signed, with negative numbers pointing to partial penetration between particles forming the two membranes, as the fusion is not allowed. Error bands correspond to the full range of values measured in all simulation replicas. Bottom: time evolution of a score assigned to the successful formation of zippered SNARE complexes in each simulation (the legend is shared between top and bottom panels). The score is calculated based on the mean pairwise distance between v-SNARE and t-SNARE particles, with higher values pointing to more successful contacts. (**b**) Distribution of membrane-membrane distance measurements for simulations including different copy numbers of curvature-inducing proteins. Distances are sampled in all simulation replicas, during the given shared time interval.

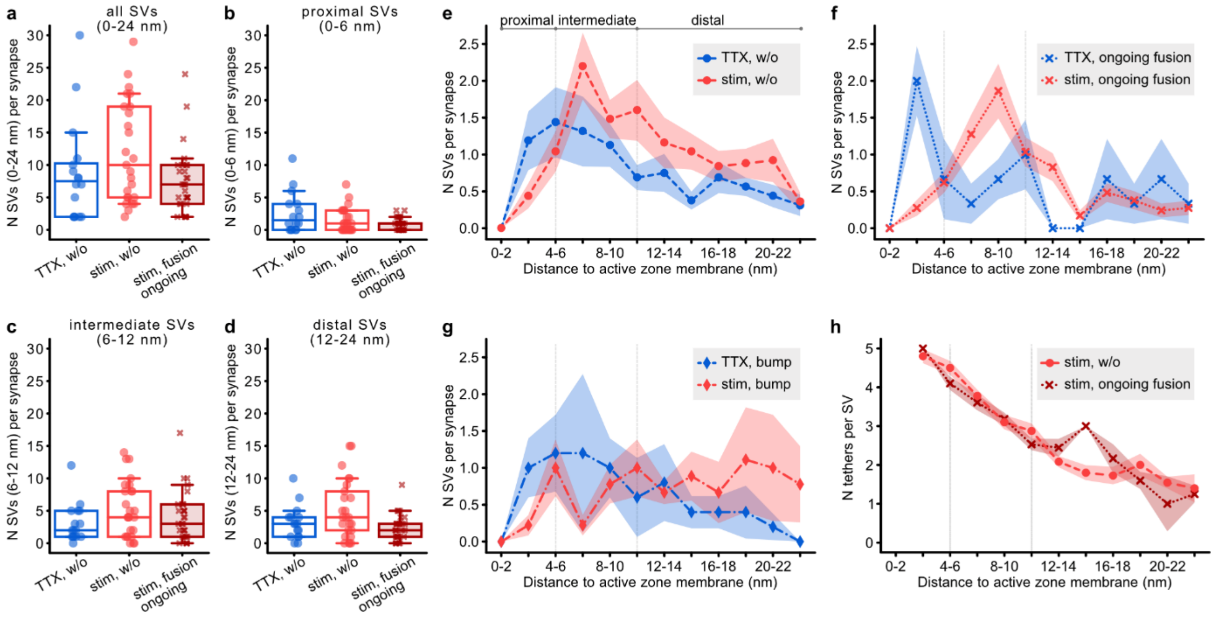


**Suppl. Fig. 9: Distributions and tethering of SVs in stimulated and control synapses.** (**a**) Number of SVs per synapse within a distance of 24 nm from the AZ membrane in TTX-treated synapses without membrane rearrangements (w/o), in stimulated synapses without membrane rearrangements, and in stimulated synapses with ongoing fusion. (**b**) Number of membrane-proximal SVs per synapse, maximal distance 6 nm. (**c**) Number of intermediate SVs per synapse, distance 6-12 nm. (**d**) Number of distal SVs per synapse, distance 12-24 nm. (**e**) Distribution of SVs in stimulated and TTX-treated synapses without membrane rearrangements. (**f**) Distribution of SVs in stimulated and TTX-treated synapses with ongoing fusion. (**g**) Distribution of SVs in stimulated and TTX-treated synapses with bumps. N TTX-treated synapses w/o = 16, N stimulated synapses w/o = 25, N stimulated synapses with ongoing fusion = 29 (for a-g). (**h**) Correlation of tether number and distance of the SV to the AZ membrane in stimulated synapses without membrane rearrangements (N = 145 SVs from 15 synapses) and in stimulated synapses with ongoing fusion (N = 114 SVs from 18 synapses).

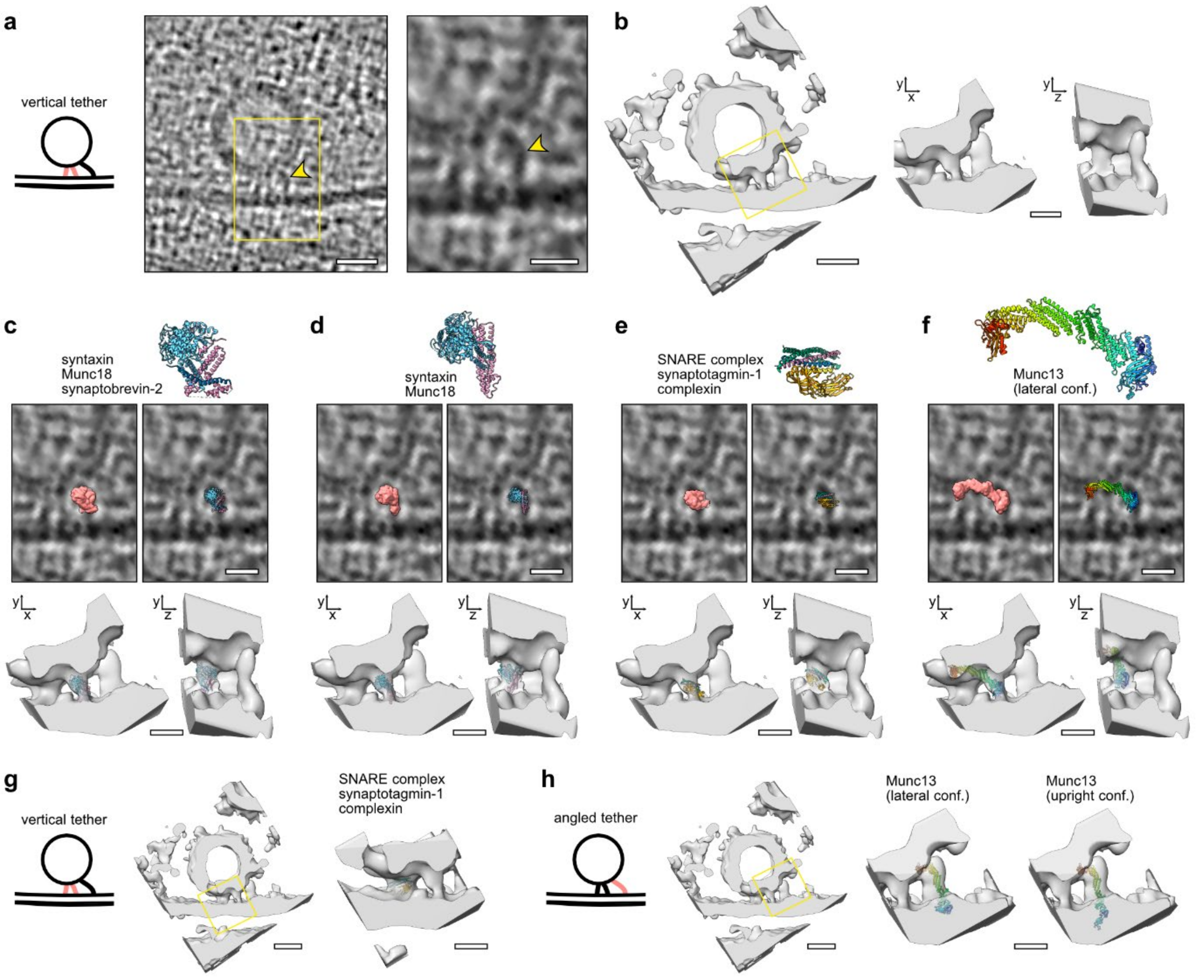


**Suppl. Fig. 10: Exemplary alignment of vesicular tether sizes and exocytic proteins.** (**a, b**) Exemplary tomogram slice (a) and isosurface (b) of a multi-tethered SV with zoom-in to tethers between SV and AZ membrane. Scale bars: whole SV 20 nm, zoom-ins 10 nm. **(c-f)** Exemplary fits of atomic models and the respective generated density maps to a vertical tether. Top row: atomic models of syntaxin/Munc18/Synaptobrevin-2 (c, PDB: 7UDB[1]), syntaxin/Munc18 (d, PDB: 4JEU[2]), an assembled SNARE complex with synaptotagmin-1 and complexin, as expected for a primed SV (e, PDB: 5W5D[3]), and Munc13 (f, PDB: 7T7V[4]). Middle row: overlays of atomic models, their respective density maps, and tomogram slices. Bottom row: fits of atomic models into isosurfaces of the vertical tether. Scale bars 10 nm. (**g**) Exemplary fit of an assembled SNARE complex with synaptotagmin-1 and complexin into the isosurface of a second, short vertical tether from the same SV. Scale bars: left 20 nm, right 10 nm. (**h**) Exemplary fit of Munc13 in its lateral (left, PDB: 7T7V) or upright conformation (right, PDB: 7T7X[4]) into the isosurface of a third, longer and angled tether from the same SV. Scale bars: left 20 nm, right 10 nm. Light blue: Munc18, pink: syntaxin-1, green: SNAP-25, dark blue: synaptobrevin-2, yellow: complexin, orange: synaptotagmin-1.

| **Category** | **Shape of the SV/vesicular part** | **AZ membrane and connection** |
|---|---|---|
| Tethered SV | Spherical SV | Flat membrane, filamentous connections (tethers) between SV and membrane |
| Tightly docked SV | Spherical SV, basal part may be flattened | Flat membrane, direct contact between SV and AZ membrane |
| Membrane invagination (Cat. 1) | Spherical SV | Slight invagination below SV, tethers between SV and membrane |
| Stalk formation (Cat. 2) | Beginning droplet formation (exvagination) | Flat membrane or slight invagination below SV, tethers and potential beginning lipid intermixing (blurry membrane) |
| Closed fusion pore (Cat. 3) | Droplet shape of the pore head, pore neck is closed | Continuous membrane (also for following categories), invagination facing the extracellular space |
| Open fusion pore (Cat. 4) | Wider droplet shape oft he pore head, pore within | Deep invagination via open pore neck, outward and inward curvature at the neck |
| Dilating fusion pore (Cat. 5) | Dome shape with vertical or angular sides | Inward curvature at the junction of SV and cell membrane (up to 90° angle) |
| Collapsing fusion pore (Cat. 6) | Flattened dome, top membrane thickened | Angular sides |
| Small bump (Cat. 7) | Slight membrane curvature, top membrane may be thickened | |

**Suppl. Table 1: Morphological characteristics of SVs and fusion states.**

### Supplementary methods

#### Electrophysiological recordings

Cell-attached voltage-clamp recordings were performed on mass cultured neurons at room temperature at days in vitro 14-15. Currents were recorded using a Multiclamp 700B amplifier (Axon Instruments) controlled by Clampex 9 software (Molecular Devices). Membrane capacitance and series resistance were compensated by 70% und data filtered by low-pass Bessel filter at 3 kHz and sampled at 10 kHz using an Axon Digidata 1322A digitizer (Molecular Devices). A fast perfusion system (SF-77B; Warner Instruments) continuously perfused the neurons with the extracellular solution which contained the following (in mM): 140 NaCl, 2.4 KCl, 10 HEPES (Merck, NJ, USA), 10 glucose (Carl Roth, Karlsruhe, Germany), 2 CaCl2 (Sigma-Aldrich, St. Louis, USA), and 4 MgCl2 (Carl Roth) (~300 mOsm; pH 7.4). The extracellular solution was supplemented with NBQX (5 µM) and bicuculine (15 µM) to block network activity. Borosilicate glass pipette containing extracellular solution were kept at resistance 2–4 MOhm, and access resistance was compensated by 70%. Optical stimulation was provided by a 470nm LED coupled into the fluorescence port of the microscope (Olympus-IX71) and triggered for 5ms by a transistor-transistor logic (TTL) signal. To evoke AP trains, 40 and 50Hz light pulse trains were triggered every 5s.

Peak amplitude and delay of the responses were calculated using Axograph X (Axograph Scientific). Delay was measured from the light onset to the peak of the evoked action potential. For each cell, 3 trials of trains were averaged to determine the final values.

#### Immunocytochemistry

After live imaging, neurons were immediately fixed with 4% paraformaldehyde in phosphate-buffered saline (PBS) for 15 min followed by a brief wash in PBS. The cells were permeabilized using 0.1% triton x-100 in PBS (PBS-T), 3 x 10 min, and fluorescence was quenched using 100 mM glycine in PBS for 10 min. Unspecific binding sites were blocked with 5% normal donkey serum in PBS-T for 1h, primary antibodies diluted in blocking solution were incubated overnight. The cells were washed 3 x 10 min with PBS-T, secondary antibodies diluted in PBS-T were applied for 1h at room temperature. Again, the cells were washed 3 x 10 min with PBS-T and once with PBS for 5 min followed by mounting with mowiol. Primary antibodies: rabbit anti-Homer1 (Synaptic systems, #160 003, dilution 1:200) and guinea pig anti-Vglut1 (Synaptic systems, #135 304, dilution 1:4,000); secondary antibodies: Rhodamine Red donkey anti-rabbit IgG (Jackson ImmunoResearch, #711-295-152, dilution 1:1000) and Alexa Fluor 647 donkey anti-guinea pig IgG (Jackson ImmunoResearch, #706-605-148, dilution 1:1000).

Confocal stacks of regions, which were imaged during iGluSnFR3 live imaging beforehand, were acquired using a Leica SP5 confocal microscope with a 63x oil immersion objective, pixel size 0.12 µm and z-steps of 0.5 µm. Overlays of live imaging and maximum intensity z-projections of confocal stacks were generated manually using fiji.

**synGCamp6f and iGluSnFR live imaging**

Room temperature live imaging of neurons (DIV14-18) infected with synGCamp6f or iGluSnFR was performed on an inverted microscope (Olympus IX71) equipped with a 60x water immersion objective, andor iXon camera (Oxford instruments), and 490 nm LED system (pE2; CoolLED). Cells were perfused with high-calcium extracellular solution containing the following: 140 mM NaCl, 2.4 mM KCl, 10 mM HEPES, 10 mM glucose, 4 mM CaCl2, and 1 mM MgCl2 (300 mOsm; pH 7.5). 6 µM NBQX and 30 µM bicuculline were added to block neuronal network activity. Action potentials (2 ms depolarization) were induced using a field stimulation chamber (Warner Instruments), Multiclamp 700B amplifier, and an Axon Digidata 1550B digitizer controlled by Clampex 10 software (all Molecular Devices). To analyze the kinetics of the biosensors, single action potentials were induced and images were captured with an exposure time of 10 ms and a frame rate of 40 fps.

**Live imaging analysis**

Live imaging recordings were analyzed using graphpad prism (room temperature) and python (elevated temperature). Stimulation and image acquisition times were extracted using axograph, fluorescence intensities of each micrograph were measured using fiji. Datasets recorded at room temperature and at elevated temperature were analyzed separately. For the analysis of synGCamp6f and iGluSnFR at room temperature, $F_0$ was calculated as the mean fluorescence intensity of the first four and the last four images of each recording. $t_{max}$ was defined as the time point at which the fluorescence intensity reached its maximum. For the calculation of $t_{on}$, a sigmoidal curve was fitted to the fluorescence intensity values between the time point of stimulation and $t_{max}$ using graphpad prism. $t_{on}$ was defined as the time point at which 50% of $t_{max}$ were reached and calculated using the sigmoidal fit. Likewise, $t_{off}$ was defined as the timepoint, at which the fluorescence intensity after stimulation decreased to 50% of its maximum, and calculated using an exponential fit. Recordings with a fluorescence intensity change ΔF/F0 of less than 0.1 were excluded from the analysis.

For the analysis of iGluSnFR constructs at elevated temperature, fluorescence intensity values were corrected for bleaching by fitting an exponential decay curve to the baseline signals before and after action potential induction. Although two APs at 40 Hz were applied, only the time point of the first AP was used for calculations. Definitions of $t_{max}$, $t_{on}$ and $t_{off}$ were the same as for the analysis of room temperature recordings. Likewise, a sigmoidal fit was used for the rise time and an exponential fit for the decay time. Again, recordings with ΔF/F0 of less than 0.1 were excluded.

**Calculation of theoretical release probability based on SV distribution**

Assuming that (1) the distribution of membrane-near SVs in synapses short after fusion resembles synapses during SV fusion, and (2) that the distribution of SVs in non-releasing synapses resembles the control condition (TTX):

(1) $N(proxSVs)_{fusion} = N(proxSVs)_{postfusion} = 0.89$

(2) $N(proxSVs)_{non\text{-}releasing} = N(proxSVs)_{TTX} = 2.54$

And (3) that the group of stimulated synapses without membrane rearrangements contains fraction x of postfusion synapses and fraction y of non-releasing synapses:

(3) $x * N(proxSVs)_{postfusion} + y * N(proxSVs)_{non\text{-}releasing} = 1.48$; with $x + y = 1$

We can calculate the theoretical fractions of postfusion synapses x = 0.83 and non-releasing synapses y = 0.17 within the group of stimulated synapses without membrane rearrangements.

Taking into account that 46% of all stimulated synapses did not show membrane rearrangements, the overall theoretical fraction of non-releasing synapses is 0.08, the theoretical release probability would thus be 0.92 based on the distribution of membrane-proximal SVs.

**Virus constructs**

| Construct | Identifier | Source/ reference |
|---|---|---|
| f(syn)ChR2(E123T/T159C)-YFP-w | BL-347 | VCF, Addgene #35511[5,6] |
| f(syn)ChR2-E123T/T159C-mscarlet-w | BL-1847 | This study |
| f(syn)ChR2-E123T/T159C-Cerulean3-w | BL-1848 | This study |
| f(syn)SynGCamp6f-w | BL-700 | VCF, Addgene #40755[7,8] |
| syn iGLuSnFR wt WPRE3 | AAV 61 | VCF, Addgene #41732[9] |
| pAAV.hSyn.iGluSnFR3.v857.PDGFR | AAV 336 | VCF, Addgene #178329[10] |
| pAAV.hSyn.iGluSnFR3.v857.GPI | AAV 337 | VCF, Addgene #178331[10] |

**Suppl. Table 2: Viral constructs.**

Virus production (lentivirus and AAV) was performed by the Viral Core Facility of the Charité – Universitätsmedizin Berlin (vcf.charite.de).

##### Subtomogram averaging of fusion states

For subtomogram averaging of fusion states, 4-times binned subtomograms (without denoising or IsoNet correction) were used. For each state, corresponding subtomograms were manually aligned in *dynamo_gallery* to produce an initial average. This was used as an initial reference for a global alignment of all subtomograms. The subtomogram average for each category was performed separately in Dynamo. Due to the low subtomogram number per class (4-10), averages were c61-symmetrized for the final representation in ChimeraX.
